# Cellular Complexity and Systemic Immune Profiles across Ancestral Diversity in Thailand and Mainland Southeast Asia

**DOI:** 10.64898/2026.06.08.726441

**Authors:** Benjamaporn Sriwilai, Sarintip Nguantad, Damita Jevapatarakul, Narita Thungsatianpun, Juthamard Chantaraamporn, Phaewa Chaiwijit, Asian Immune Diversity Atlas Network, Ankita Chatterjee, Partha P. Majumder, Jay W. Shin, Yoshinari Ando, Jong-Eun Park, Woong-Yang Park, Kian Hong Kock, Le Min Tan, Shyam Prabhakar, Manop Pithukpakorn, Bhoom Suktitipat, Ponpan Matangkasombut, Varodom Charoensawan

## Abstract

Mainland Southeast Asia (MSEA) remains under-represented in global immunogenomic references despite its extensive genetic heterogeneity. We present the first single-cell immune atlas of an MSEA population, utilizing Thai individuals from the Asian Immune Diversity Atlas (AIDA) as a representative cohort. We demonstrate that the Thai population is highly genetically diverse, reflecting its history as a geographic nexus for Asian admixture. By integrating single-cell transcriptomics with high-resolution genotyping, we show that genetic ancestry significantly shapes innate immune profiles, specifically CD14^+^ monocytes, highlighting potential evolutionary adaptations to regional pathogens. We identify specific sex-by-ancestry interactions that may drive the baseline activation of pro-inflammatory pathways in females, providing a long-sought cellular rationale for the high prevalence of autoimmune disorders observed in Southeast Asian populations. Ultimately, our study reveals that population-specific genetic architecture dictates immune heterogeneity often missed by self-reported ethnicity or country of origin, providing a critical immunogenomic reference for precision medicine in MSEA regions.

## INTRODUCTION

Mainland Southeast Asia (MSEA) possesses one of the most complex demographic histories since the initial “Out of Africa” migration, shaped by millennia of successive admixture events. Consequently, the current MSEA populations represent a nexus of diverse ethnicities, remaining historically, culturally, and geographically distinct from Island Southeast Asia (ISEA) ^1-3^. Situated at the geographic center of the region, Thailand shares deep historical and environmental connections with surrounding MSEA nations, namely Myanmar, Laos, Cambodia, Vietnam, and Malaysia, making it highly representative for the region. Historical migrations and series of environmental pressures, including a wide range of tropical pathogens, have imprinted signatures of natural selection upon the MSEA genome in terms of immune-related traits ^4^. However, genomic data alone are not sufficient to elucidate how genetic variations dictate cellular heterogeneity and functional immune diversity. Interplay between genetic ancestry and the immune profiles of contemporary MSEA populations remains poorly characterized.

Previous studies of diverse populations have established that genetic ancestry is a primary determinant of immune response variation. For instance, ancestry-specific polymorphisms affecting *TLR1* expression in Europeans are associated with reduced pro-inflammatory monocyte response to bacterial challenges, relative to individuals of African descent ^5^. Similarly, monocyte-derived macrophages from individuals of African ancestry exhibit stronger bacterial control compared to those of European ancestry ^6^. Interestingly, divergent human leukocyte antigen (HLA) expression patterns linked to African ancestry have also been implicated in differential susceptibility to autoimmune conditions, including rheumatoid arthritis and ulcerative colitis ^6^. More recent single-cell studies further demonstrate that ancestry-driven signatures are highly cell-type specific, notably, divergent monocyte transcriptional profiles in European and African cohorts have been linked to differential susceptibility to influenza A ^7^ and varying degrees of COVID-19 severity ^8^.

To date, single-cell transcriptomic immune profiling has largely been restricted to European cohorts, e.g., ^9,10^. Similar efforts are only beginning to emerge for Asian populations, despite representing the largest global demographic. Recent immune profiling efforts involving Asian populations and their descendants include comparative analyses of immune characteristics between healthy individuals and those diagnosed with autoimmune diseases across both European and Asian ancestries, such as the California Lupus Epidemiological Study (CLUES), the Immune Variation Project (ImmVar) cohorts ^11^, and the ImmunoMicrobiome cohort ^12^. More recently, the Asian Immune Diversity Atlas (AIDA) provides a single-cell reference of over 1 million immune cells from more than 600 donors across five Asian countries, namely Japan, South Korea, Singapore, India, and Thailand, revealing how sub-continental ancestry, age, and sex drive cellular and molecular variation ^13^. Furthermore, the Chinese Immune Multi-Omics Atlas (CIMA) offers a high-resolution map integrating transcriptomic and epigenomic data from more than 400 individuals ^14^. While these resources address the historical bias toward European ancestry, they often rely on self-reported ethnicity or focus on relatively homogeneous East Asian groups or Western-based Asian diaspora. Such approaches may overlook the distinct genetic complexity and ancestral signatures inherent to Southeast Asian populations residing in their Mainland and Island Southeast Asia environments. Only recently has the immune landscape of ISEA been characterized, specifically in 199 individuals from Bali and New Guinea ^3^. Consequently, the relationship between fine-scale genetic admixture and immune variation in contemporary MSEA populations remains insufficiently characterized.

To address this gap, we present single-cell immune characterization of 59 contemporary Thai individuals, as representative of the MSEA population. Building on the first single-cell immune profiles of Thai individuals as part of the AIDA cohort, we integrate high-resolution genetic variation information with single-cell transcriptomics, to elucidate how specific ancestral lineages, distinct from the East and South Asian populations, contribute to cellular heterogeneity and immune diversity. In addition to the ancestry components, we utilize multiple linear regression (MLR) and differential abundance (DA) analyses to assess the impact of sex and age on immune cell proportions and their differential gene expression profiles, taking into account these covariate factors within a unified modeling framework. Our work establishes a foundational resource for MSEA and provides the necessary resolution to facilitate future disease-association studies tailored to the unique admixed genomic architecture and distinct environmental exposures of this melting pot region.

## RESULTS

### The first representative immune profiles of present-day Thai and MSEA populations

We first characterized the genotypes of 59 representative healthy Thai individuals, with single-cell immune profiles available through the Asian Immune Diversity Atlas (AIDA) consortium ^13^, by contextualizing them against both modern and ancient individuals from major Asian and global populations in the Allen Ancient DNA Resource (AADR) ^15^. Overall, genomic profiles of the Thai and neighboring MSEA populations are highly diverse and positioned at the intersection of East Asian (EAS), South Asian (SAS), and Island Southeast Asian (ISEA) ancestries **(Figure 1A and S1)**, consistent with their geographical distribution in the analysis (**Figure 1B)**. Importantly, the genomic profiles of these representative Thai individuals are the most closely aligned with the Mainland Southeast Asian (MSEA) populations from the AADR, particularly those from Myanmar, Laos, Cambodia, Vietnam, Malaysia **(Figure 1A)**, as well as minority subgroups within Thailand, including Sgaw Karen, Lawa, Tai Lue, Akha, Hmong, Htin Mal, Mlabri, Mon, Nyah Kur, Khmer, Kuy Suay, and Maniq (**Figure S1)**. This suggests that the Thai and MSEA populations share similar genetic structures, and the single-cell immune profiles from 59 individuals in the AIDA Thailand cohort also serve as a proxy for the immune profiles of broader MSEA populations and associated ethnic groups under shared environmental and selective pressures in the tropical region.

**Figure 1.**
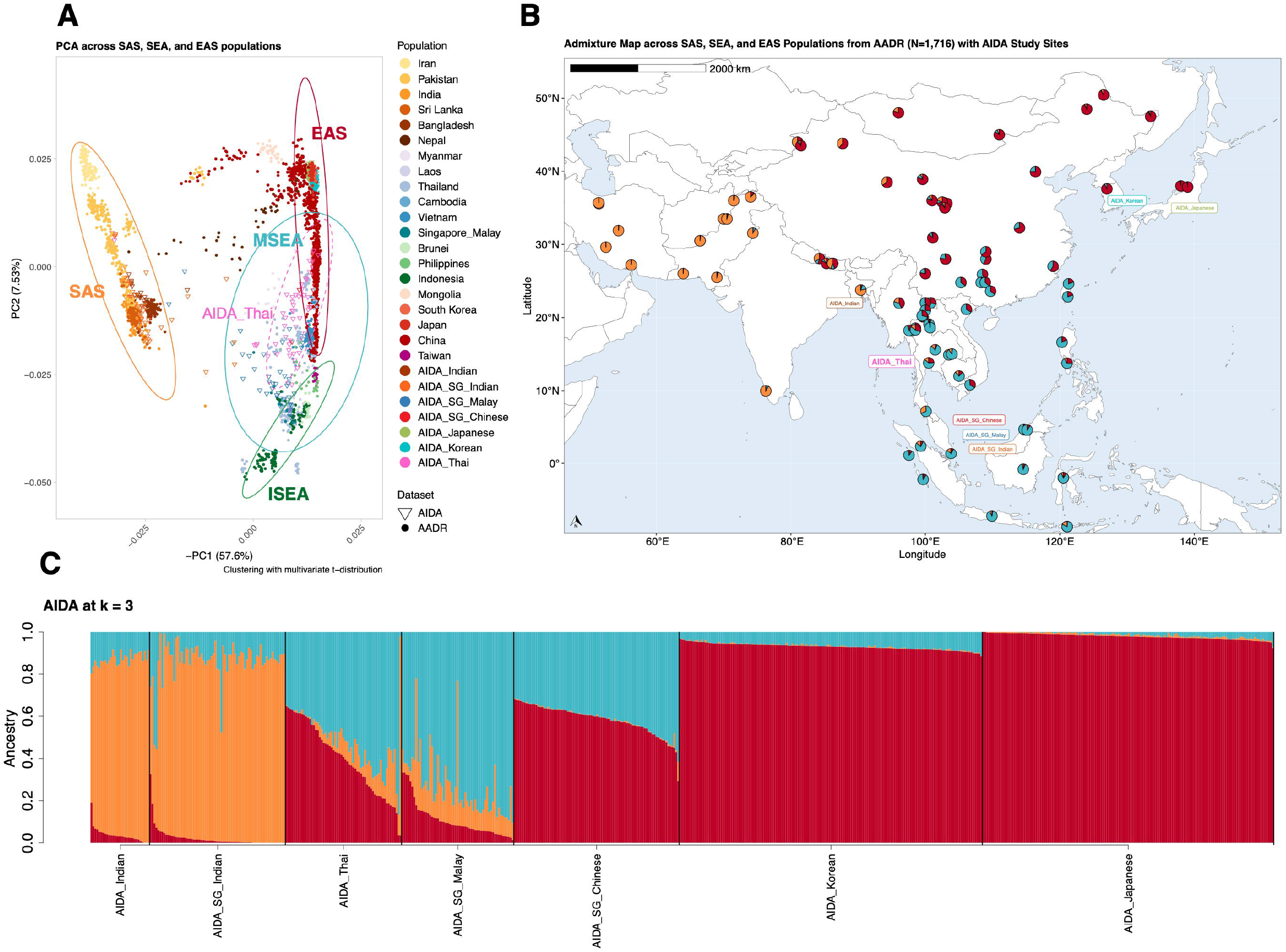
Present-day Thai is genetically diverse. **(A)** Principal Component Analysis (PCA) illustrates genetic variation among 2,734 unrelated samples, including AIDA samples and reference populations from East Asia (EAS), South Asia (SAS), and Southeast Asia (SEA). **(B)** Admixture map showing ancestry proportions across populations, overlaid on their geographical locations as a representation from AIDA and AADR samples. **(C)** Ancestral components per individual inferred from genotypes of 601 unrelated AIDA samples using ADMIXTURE at k = 3. Genetic ancestry estimation was performed after imputation and quality control on 113,832 autosomal SNPs of 2,734 unrelated samples.

To investigate genetic diversity beyond the resolution of countries of recruitment, cohorts, or self-reported ethnic groups, we examined the ancestral components of the AIDA cohorts and Asian populations in the AADR using ADMIXTURE analysis **(Figures 1B and 1C)**. Three major ancestral components were identified, each represented by an anchoring population: (1) Orange - predominant in Southwest and South Asian populations (with AADR’s “ancient Iranians” as an anchoring population); (2) Red - representing the primary ancestral component associated with of in East Asia populations (“Qiang” as an anchoring population); (3) Blue - characteristic of nomadic hunter gatherers from non-Negrito groups (“Mlabri” as an anchoring population), who are closely related to Hoabinhians, an ancient lineage which contributes substantially to present-day Mainland Southeast Asian and Island Southeast Asian populations ^2,16^. Cross-validation error estimates were robust to differences in sample sizes **(Figures S2A and S2B)** and variant sets **(Figure S3A and S3B)**.

Focusing on the ancestral composition of the representative Thai AIDA cohort and other MSEA populations, all three major components were observed, with a broad gradient of admixture primarily between the Red and Blue ancestries, and a smaller contribution from the Orange component. This pattern further supports the view of MSEA as a genetic and cultural crossroads, reflecting its historical role as a central intersection among EAS, SAS, and ISEA (**Figure 1C**). We further quantified the genetic diversity using Shannon entropy and found that the representative Thai cohort exhibited the highest degree genetic diversity resulting from high levels of admixture from all the three ancestral components, as compared to other AIDA cohorts (**Figure 1C**) and most Asian populations in the AADR is comparable to that of MSEA populations such as Burmese and Thai Mon (see **Table S1)**.

### Comprehensive evaluation of the effects of self-reported ethnicity, age, and sex on immune cell composition in the AIDA cohorts

Having characterized the genomic backgrounds of representative Thai individuals in the context of MSEA and other major Asian populations, we next examined their immune profiles, leveraging the comprehensive single-cell peripheral blood mononuclear cell (PBMC) resource generated by the AIDA consortium. We first examined the 35 immune cell subtypes annotated and described in the AIDA main publication ^13^, “AIDA marker paper” herein **(Figures 2A and S4)**, and investigated how ancestry influences immune cell compositions, while accounting for potential confounding effects of age, sex, and recruitment sites.

**Figure 2.**
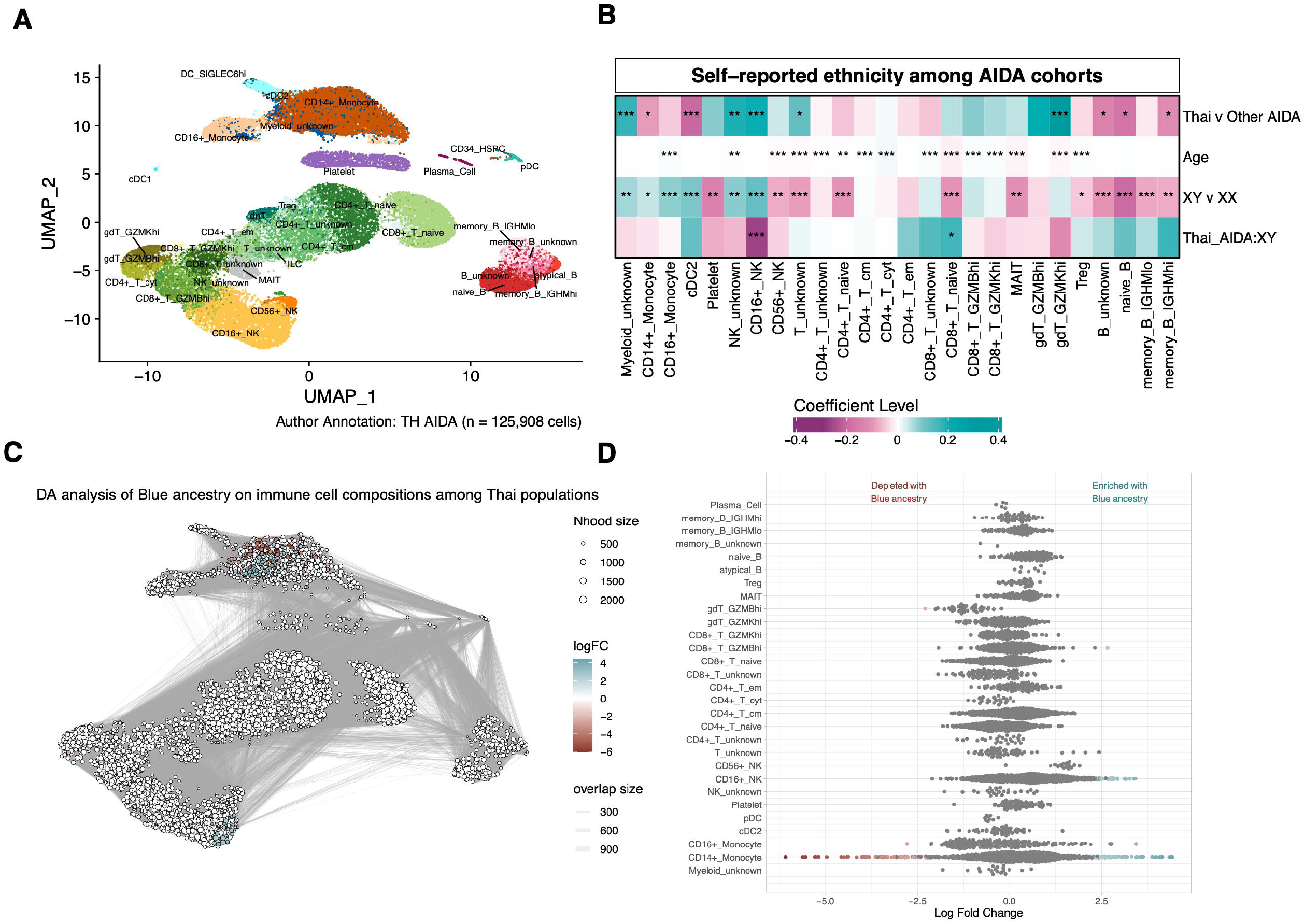
Immune cell types among Thai populations. **(A)** Uniform Manifold Approximation and Projection (UMAP) showing immune cell types in Thai donors. **(B)** Correlation of self-reported ethnicity, age, and sex on immune cell proportion between the Thai cohort and other AIDA populations (log10(Proportion) ∼ Cohort + Age + Sex + Cohort:Sex). Statistical analyses were performed using MLR; * *p* < 0.05, ** *p* ≤ 0.01, *** *p* ≤ 0.001. **(C, D)** The association of Blue ancestry on immune cell abundances among Thai populations. The statistical testing was using GLMM for each neighborhood (Cell count_nhood_ ∼ Age + Sex + % Blue ancestry + (1|Library) at spatial FDR < 0.05).

As first observed in the AIDA marker publication, the Thai AIDA cohort exhibited certain distinctive immune cell-type compositions as compared with the rest of other self-reported ethnicities within the AIDA. Notably, the Thai cohort showed lower proportions of monocytes and cDC2 conventional dendritic cells, as well as a lower proportion of CD16^+^ natural killer (NK) cells in the XY compared with XX individuals ^13^. Here,we systematically reassessed the differences in cell-type compositions between the Thai and other AIDA cohorts using multiple linear regression (MLR), incorporating age and sex as covariates, and the interaction terms between age, sex, and the Thai AIDA cohort (see **Methods and Figure S5A**).

Our MLR model recapitulated age and sex-associated relationships of cell-type compositions that were consistent across the Thai and other AIDA cohorts, including the increased proportions of CD16^+^ monocyte, central memory CD4^+^ T cells, cytotoxic CD4^+^ T cells, CD8^+^ T cells with high expression of granzyme genes, and regulatory T cells (Treg) **(Figures 2B and S5A)**. In contrast, CD56^+^ NK cells, CD8^+^ T naïve cells, Mucosal-associated invariant T (MAIT) cells, and γδ T cells with high granzyme K expression were progressively depleted with age. Sex-biased differences in immune cell proportions towards XY individuals were observed for myeloid innate cells (i.e., monocytes and cDC2); whereas several lymphoid populations (i.e., CD56^+^ NK, naive T cells, MAIT, Treg, and B cells) were enriched in XX individuals. Interestingly, this age- and sex-adjusted MLR model also revealed the cohort-specific interactions of the Thai cohort and sex differences. Proportions of CD16^+^ NK cells were more pronounced in XY individuals across all AIDA self-reported cohorts, except the Thai cohort, where they were more enriched in XX. A similar reversal of sex-associated differences was observed for CD8^+^ T naive proportion between the Thai and other AIDA cohorts **(Figure 2B)**.

Regarding differences in cell-type proportions between the Thai and other AIDA populations based on self-reported ethnicity, our models revealed consistent findings with the previous analysis, showing reduced proportions of CD14^+^ monocytes and cDC2 in the Thai cohort ^13^. Apart from these innate myeloid populations, we also identified differences across lymphoid lineage cells. Specifically, the Thai cohort exhibited lower proportions of some B cell subtypes and higher proportions of some NK and γδ T cells compared with other AIDA cohorts **(Figure 2B)**. However, these differences remain inconclusive and may reflect potential confounding by genetic backgrounds and recruitment sites.

### Dissecting the impact of admixed ancestry on immune profiles in Thai and other Asian populations

As established in the first section, MSEA and other Asian populations are highly admixed and comprise overlapping ancestral components present in varying proportions. Consequently, the impact of genetic diversity on immune profiles cannot be reliably assessed from the country of recruitment or self-reported ethnicity alone. To address this limitation, we re-performed the MLR of cell type abundances, incorporating the three inferred ancestral components as predictors in place of self-reported ethnicity of all healthy individuals in the AIDA **(Figure S5B)**, as well as of each individual AIDA cohort **(Figure S6A)**. Overall, this ancestry-associated MLR recapitulated most of the age- and sex-associated differences in immune cell proportions observed using the self-reported ethnicity model, but also revealed ancestry-specific associations, including the enrichment of Treg and naïve B cell populations with increasing Orange ancestry, and their relative depletion towards the Red ancestry in the Thai populations **(Figure S6B)**.

Next, we investigated the influence of ancestry on subpopulation of cells with distinct gene expression profiles within major immune cell types using differential abundance (DA) analysis, while correcting for technical confounders such as experimental batch effects. Although the variables of interest, namely ancestral components, age, and sex, were evenly distributed across the four batches of the Thai AIDA cohort **(Figure S7)**, all were nonetheless included as covariates in the DA models to ensure robust inference. Consistent with the previously mentioned age-associated increase in overall CD16^+^ monocyte proportions and CD8^+^ T with high *GZMB* expression **(Figure 2B)**, we further identified specific subpopulations that were significantly enriched with age in the Thai and other AIDA cohorts **(Figure S8)**. In addition, although CD16^+^ NK cells did not exhibit an age-associated effect in the MLR model of overall cell-type proportions, yet the DA analysis revealed specific subpopulations that were enriched with age. In contrast, the proportions of naive CD8^+^ T and MAIT cells decreased with age, along with age-associated depletion of their respective subpopulations. Regarding sex-specific immune subpopulations, CD16^+^ NK cells were predominantly enriched in XY individuals across most AIDA populations but showed greater enrichment in XX individuals specifically within the Thai cohort (**Figure S9**). In contrast, naive CD8^+^ T cells were enriched in XY individuals specifically in the Thai cohort. Other sex-associated subpopulations include myeloid cells, e.g., CD16^+^ and CD14^+^ monocytes and cDC2, displayed enrichment patterns that were unique or shared between the Thai cohort and several other AIDA populations.

Focusing on the influence of ancestral components within the Thai AIDA cohort, we identified enrichment of specific CD14^+^ monocyte and CD16^+^ NK subpopulations associated with increased Blue ancestry (**Figures 2C, 2D and** see **Table S2)**. The Blue component represents descendants of Hoabinhian hunter-gatherers, who inhabited the region prior to successive waves of East Asian farmer migrations, which is likely represented by the Red ancestry. Notably, these associations were evident at the subpopulation level and were not detectable when analyzing the corresponding cell types in aggregate (**Figure S6A)**. Such ancestry-associated innate immune subpopulations were not observed in the Singaporean Chinese (SG Chinese) or Singaporean Malay (SG Malay) cohorts when analyzed separately, despite both groups comprising predominantly Red and Blue ancestral components and having comparable sample sizes to the Thai cohort **(Figure S10B)**. However, when the three SG cohorts were combined prior to analysis, thereby recapitulating the broader spectrum of admixture observed in the Thai cohort, the Blue ancestry-associated CD14^+^ monocyte and CD16^+^ NK subpopulations became detectable, together with enrichment of certain innate-like T cells (i.e., circulating γδ T cells with high expression of granzyme B and MAIT cells) **(Figure S10C)**. Together, these findings suggest that genetic admixture contributes to shaping variation within the innate immune system, and also indicate that analyses based solely on overall cell-type proportions lack sufficient resolution to detect ancestry-associated immune heterogeneity at the subpopulation level.

### Genetic ancestry diversity underlines distinct innate immune responses associated with pro-inflammatory and interferon pathways

Building on the ancestry-associated CD14^+^ monocyte and CD16^+^ NK subpopulations observed in the Thai and combined SG AIDA cohorts described earlier, we expanded the DA analysis to investigate genes associated with ancestry-associated subpopulations of myeloid innate immune cells (i.e., monocytes, cDC, dendritic cells), as well as innate lymphoid cells (ILCs) and NK cells at the higher resolution of cell-type specific transcriptome profiles (see **Methods**). This more fine-grained DA analysis confirmed the presence of Blue ancestry-associated subpopulations of CD14^+^ monocytes (**Figures 3A and 3B**) and CD16^+^ NK cells (**Figures S11A and S11B**) in the Thai cohort, and this finding was not attributable to batch effects due to cell type identification and demographic distribution **(Figures 3A, S11A, S12A, S13A, and S7)**. Again, these Blue ancestry-associated subpopulations were not detected in other AIDA cohorts defined by self-reported ethnicity, but became evident when the three SG self-reported cohorts were combined **(Figure S14)**.

**Figure 3.**
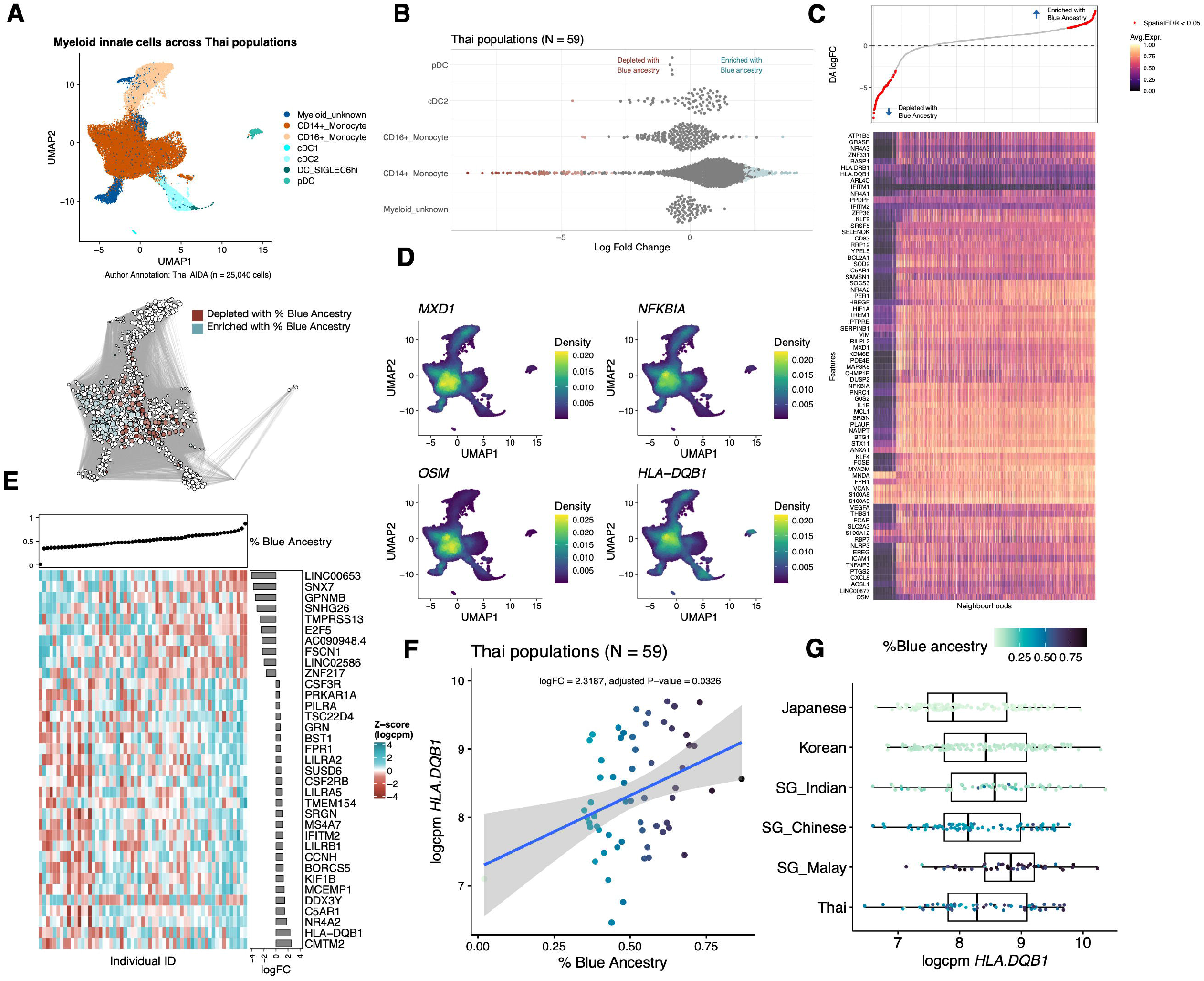
Association of the Blue ancestral component with cell states and gene expression profiles of myeloid innate immune cells in the Thai populations. **(A)** UMAP and neighborhood enrichment of myeloid innate cells associated with Blue ancestry in Thai populations, assessed using GLMM per neighborhood (Cell count_nhood_ ∼ Age + Sex + % Blue ancestry + (1|Library), spatial FDR < 0.05). **(B)** Beeswarm plot showing association between Blue ancestry and differential neighborhoods abundance across major cell types. **(C)** Representative up-regulated genes in CD14^+^ monocyte neighborhoods correlate with increasing Blue ancestry (MLR: Gene expression_nhood_ ∼ Age + Sex + % Blue ancestry at logFC > 0.5 and adj.P.Val < 0.05). **(D)** Density plots of representative genes up-regulated by increasing Blue ancestry using Nebulosa v1.0.1 ^47^. **(E)** DEGs in CD14^+^ monocytes associated with Blue ancestry, based on Pseudobulk transcriptome per individual using an LMM at |logFC| > 0.5 and adj.P.Val < 0.05. **(F)** The correlation between Blue ancestry component and *HLA-DQB1* expression in CD14^+^ monocytes across 59 Thai individuals and **(G)** in comparison with other AIDA populations. For visualization of **(E-G)**, gene expression was aggregated across two libraries per individual and normalized using TMM across all genes.

Focusing on the ancestry-associated neighborhoods of CD14^+^ monocyte in the representative Thai healthy individuals, we observed high expression of interferon-stimulated genes (e.g., *MXD1, IFITM1, IFITM2*), cytokine-mediated immune and inflammatory mediators (e.g., *NFKBIA, OSM, IL1B*, and *TNFAIP3*), as well as HLA genes (e.g., *HLA-DQB1, HLA-DRB1*) in the neighborhoods associated with high proportions of Blue ancestry (**Figure 3C**, example up-regulated genes in **Figure 3D; Table S3**). Similarly, when all the three SG cohorts were analyzed together, increasing Blue ancestry was again associated with enrichment of interferon-stimulated genes within CD14^+^ monocyte neighborhood (e.g., *GBP1, GBP4, GBP5, IFITM2*), (**Figure S15 and Table S4)**. This is consistent with the up-regulated genes reported in SG Malay individuals in the AIDA marker publication ^13^, where SG Malay was analyzed as a self-reported ethnic group. Genes highly expressed in the Blue ancestry-associated CD14^+^ monocyte neighborhoods are related with pro-inflammatory immune signaling pathways and interferon gamma responses in the Thai cohort, while the combined SG cohorts present association with only interferon response pathways **(Figure S16 and Table S5)**. The other ancestry-associated immune cell type in the Thai populations, CD16^+^ NK cells, exhibited a similar pattern, with neighborhoods enriched for higher proportions of Blue ancestry also showing strong associations with interferon response pathways (e.g., *IFITM1* and *IFITM2*) (**Figures S11C and S11D**; **Table S6**).

If ancestry is to serve as a meaningful predictor of an individual’s immune cell state, gene expression profiles must be analyzed at the individual level rather than at the cell neighborhood level. Consequently, we investigated genes whose pseudobulk expression profile of each individual in the Thai cohort were most strongly associated with variation in the Blue ancestry component **(Figure 3E**). As seen in the DA analysis described above, individual-level expression of several genes in interferon response pathways (e.g., *IFITM2, HLA-DQB1, C5AR1, LILRB1*) demonstrated up-regulation with increasing Blue ancestry in CD14^+^ monocytes (**Figure 3E and Table S7)**. This pattern was not observed in CD14^+^ monocytes from any other AIDA population if analyzed as separate cohorts, but if the three Singaporean cohorts were combined, elevated expression of several pro-inflammatory genes (e.g., *HLA-DQB1, TNF, MARCO, IFITM3)* became more apparent **(Figures S17 and S18)**. This finding is further supported by the enrichment of genes associated with Blue ancestry in both Thai and combined Singapore cohorts for interferon gamma and alpha response pathway **(Figure S19)**. Finally, we examined whether this phenomenon also persisted when the entire AIDA population was analyzed as a single cohort. Indeed, we observed positive correlations between increasing Blue ancestral component and pseudobulk gene expression in CD14^+^monocytes of interferon-induced genes (e.g., *GBP1*) and *HLA* genes of MHC class II in CD14^+^ monocytes (i.e., *HLA-DRB1, HLA-DQB*1, *HLA-DRA*) **(Table S8)**.

### Impact of genetic ancestry, sex, and age on differential gene expression across Thai and AIDA cohorts

Having examined the relationships between individual-level pseudobulk gene expression and Blue ancestry component in CD14^+^ monocytes **(Figures 3E-3G)**, we extended similar differential expression (DE) analysis to other immune cell types (see **Methods). Figure 4A** confirms that the number of differentially expressed genes (DEGs) up-regulated with increasing Blue ancestry is highest in CD14^+^ monocytes (10 down-regulated and 15 up-regulated DEGs with increasing Blue ancestry, using logFC>1 and adj p-value <0.05) while the number of DEGs down-regulated with increasing Blue ancestry is highest in CD16^+^ NK cells (49 down-regulated and 3 up-regulated DEGs, using the same thresholds) in the representative Thai cohort. Consistent with earlier DA analysis, DEGs positively correlated with Blue ancestry in CD14^+^ monocytes include genes involving antigen presentation, inflammation, and interferon responses (e.g., *HLA-DQB1, IFITM2, C5AR1, LILRB1*) (**Figures 3F, 3G and S20)**. Conversely, DEGs down-regulated with increasing Blue ancestry include *ZNF217, GPNMB*, and *E2F5*, which are key regulators of cell proliferation, maturation cell differentiation, and cell cycle progression **(Table S7 and Figure S20)**. Interestingly, we also observed a few DEGs in adaptive immune cells, even though an association between Blue ancestry and adaptive immune cells was not detected by the ancestry-associated MLR nor DA analyses described earlier (**Figures 4A and S21**). Blue ancestry-associated genes in adaptive immune cells include *CENPK*, which is involved in cell mitosis and shows higher expression levels in those with high Blue ancestry component, in central memory CD4^+^ T cells and naive CD8^+^ T cells. Up-regulation of *HLA-DQB1* with increasing Blue ancestry was also found in other antigen presenting cells (i.e., switched memory B cells) (**Table S7 and Figure S21)**. We then asked whether similar associations between Blue ancestry-related DEGs and specific immune cell types can be observed in the combined SG cohorts, which represent a similar broad spectrum of admixture observed in the Thai cohort. Indeed, the highest numbers of DEGs with Blue ancestry were observed in innate-like T cells: MAIT and γδ T cells (**Figure S22A**), consistent with the DA described in the previous section **(Figure S10C)**. For example, *HLA-DRB1* was up-regulated with Blue ancestry in γδ T cells, while the down-regulation of *KLRD1* in MAIT cells and *EGR1* in γδ T cells, was observed with increasing Blue ancestry among in the combined SG cohorts **(Figures S22B and S22C)**.

**Figure 4.**
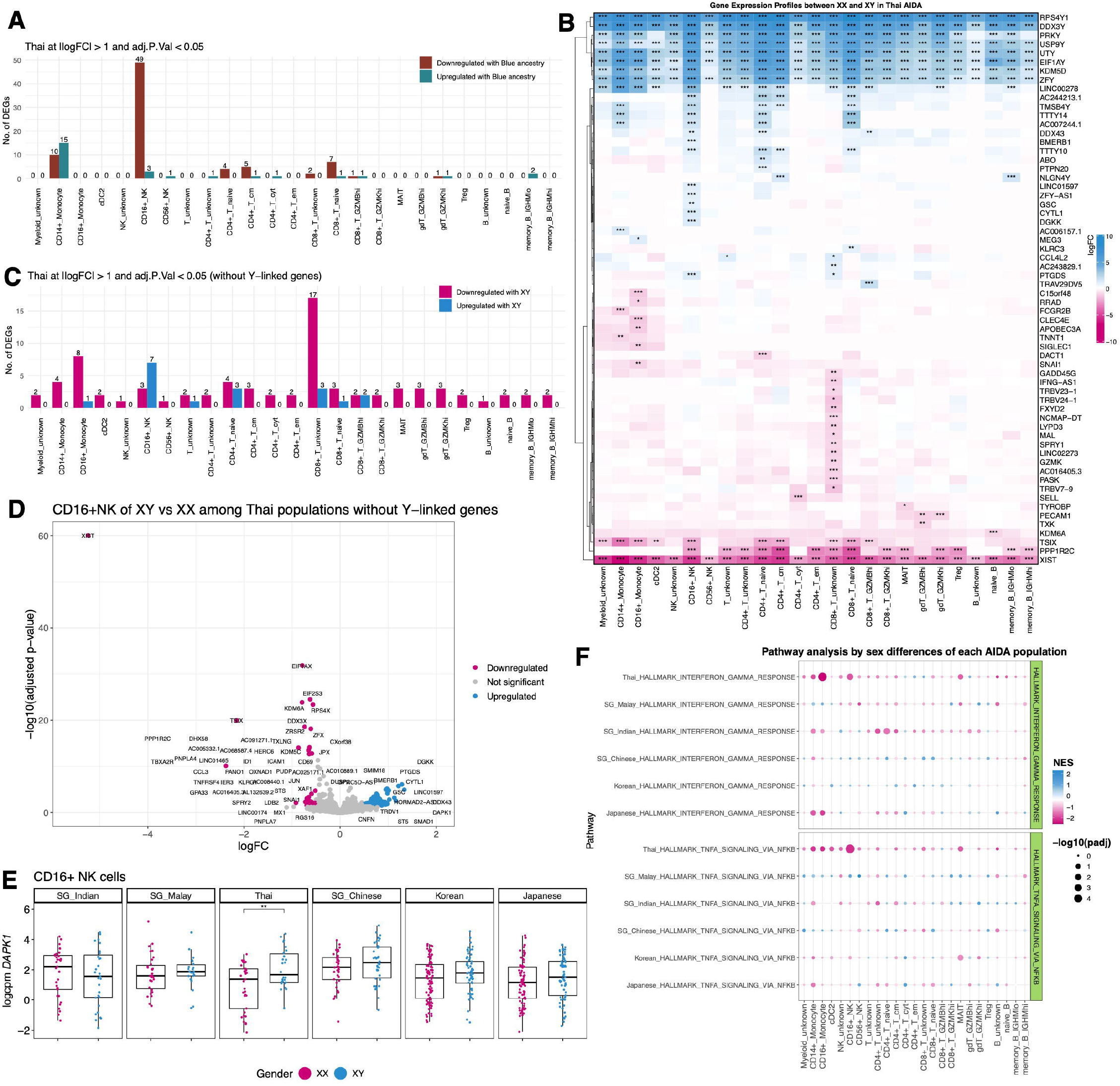
Association of Blue ancestral component and sex differences with immune profiles in Thai populations. **(A)** Number of DEGs associated with Blue ancestry in each immune cell type. **(B)** DEGs associated with sex in Thai populations; * |logFC| > 1 and adj.P.Val < 0.05, ** |logFC| > 1 and adj.P.Val ≤ 0.01, *** |logFC| > 1 and adj.P.Val ≤ 0.001. **(C)** Number of DEGs derived from X-linked versus autosomal genes. **(D)** Volcano plot showing sex-associated transcriptomic differences in CD16^+^ NK cells. **(E)** *DAPK1* expression in XY and XX individuals in the Thai cohort compared with other AIDA populations. **(F)** Top enriched pathways in the Thai population compared with other populations.

In addition to the ancestry component, we next examined the associations of gender and age with gene expression across major immune cell types in the Thai and AIDA individuals, following similar DEG analysis described above. As we observed higher abundances of CD16^+^ NK cells in XX and naive CD8^+^ T cells in XY specifically in the Thai cohort **(Figure 2B and S9)**, we next examined sex-associated DEGs to explore their potential functional roles and biological implications. We observe distinct expression patterns of both sex-chromosome genes and autosomal genes between XX and XY individuals across Thai (**Figure 4B**) and other AIDA populations **(Figure S23)**. After excluding Y-chromosome genes (**Figure 4C and S24**), the most pronounced expression differences between XX and XY were observed for X-linked genes, e.g., *XIST, TSIX, PPP1R2C, KDM6A*. These XX-associated DEGs were detected across most of the immune cell type analyses, reflecting a broad, cell-type-independent signature of sex-associated transcriptional variation (**Figure S23**). Interestingly, not all the X-chromosome DEGs showed higher expression in XX individuals. For instance, *DGKK*, which regulates lipid signaling by converting diacylglycerol to phosphatidic acid, was up-regulated in CD16^+^ NK of XY individuals compared to XX, in Thai and other AIDA cohorts **(Figure 4B and S23)**. Beyond the sex-chromosome genes in CD16^+^ NK cells, we also identified a unique up-regulation of *DAPK1*, a mediator of gamma-interferon-induced programmed cell death of the Thai XY as compared to XX individuals. A similar trend was observed in the Japanese, Korean, and SG Chinese cohorts, but the difference was most pronounced in the Thai AIDA cohort **(Figures 4D and 4E)**. In contrast, no individual autosomal genes were significantly down-regulated in Thai XY compared to XX individuals, when applying the same thresholds (logFC>1 and adj p-value <0.05) (**Figures 4C and 4D**). However, GSEA provided additional resolution, revealing a significant depletion of the TNF signaling pathway via NF-κB in the Thai XY group **(Figures 4F and S25; Table S9)**. This enrichment was driven by a subset of genes, notably *TNFRSF4, CD69, JUN*, representing the most pronounced differences in this comparison (**Figure 4D and Table S9**). In naive CD8^+^ T cells, which were specifically more abundant in the Thai XY than XX individuals, we observed up-regulation in XY of *KLRC3*, a key gene involved in HLA-E recognition **(Figure 4B, S23, and S26)**.

Other immune cell types that demonstrated unique gender-biased characteristics include CD16^+^ and CD14^+^ monocytes, and cDC2 in the Thai AIDA population (**Figure 4B and 4C**), although our MLR and DA analyses showed gender-biased cell abundances and subpopulations shared by several AIDA cohorts (**Figure S9**). This unique sex-biased down-regulation was observed exclusively in the Thai XY individuals, characterized by the down-regulation of genes involved in interferon-gamma signaling, such as *SIGLEC1* of CD16^+^ monocytes compared to XX Thai populations **(Figure 4B, S23, and S26; Table S9)**. Finally, we investigated the influence of ages on DEGs using the same thresholds (logFC>1 and adj p-value<0.05) but did not detect significant age-associated DEGs. This likely reflects the more subtle changes of expression across the age range (max absolute logFC<0.1), as compared to more pronounced effects of ancestry and sex, which were readily detectable under these criteria of every AIDA cohort.

## DISCUSSION

In this study, we present the first high-resolution single-cell immune atlas of a contemporary Thai cohort, providing a critical reference for the ancestrally and culturally complex MSEA population. Our findings reveal that the contemporary Thai population is genetically diverse as compared to other AIDA cohorts, mirroring its geographic position at the intersection of South, East, and Islands Southeast Asia (ISEA). Unlike the more homogeneous groups often captured in continental-scale studies, the Thai cohort exhibits distinct ancestral patterns that diverge significantly from both East Asian and ISEA populations. These signatures align with complex demographic shifts spanning from the Last Glacial Maximum to the Neolithic transition, introducing multiple waves of admixture and shaping present-day Thai and MSEA genomes^1,2,17,18^.

The extensive genetic admixture within the Thai population makes it a valuable resource for studying the intersection of ancestry and immunity. This diversity effectively captures the ancestral complexity of the broader MSEA region within a single cohort, potentially providing a high-resolution window into regional immune regulation. Notably, certain ancestry-associated immune characteristics that remain undetectable in other AIDA populations, including each of the three SG population groups individually and these signals only emerged when the three SG cohorts were combined to represent a broader range of genetic diversity. Consequently, Thailand serves not only as a geographic center, but potentially as a biological accelerator for immunogenomic discovery, offering the necessary resolution to dissect how complex lineage mixing dictates cellular heterogeneity and functional immune variation.

Consistent with recent genomic evidence indicating that immune-related loci in MSEA populations have undergone positive selection and pathogen-driven adaptation ^4^, our single-cell transcriptomic data demonstrated how these evolutionary signatures manifest as distinct cellular phenotypes. Moving beyond self-reported ethnicity, we leveraged high-resolution genomic data from SNP arrays, initially intended for donor demultiplexing within the multiplexed single-cell library preparation applied across the AIDA cohorts ^13^, to resolve fine-scale genetic structure. This approach allowed us to surpass broad categorical labels and instead, investigate immune variations as a function of continuous ancestral gradients. In addition to the two main lineages, Red (EAS-related ancestry) and Orange (SAS-related ancestry), within the AIDA populations, we identified the Blue ancestry component, which likely descends from the pre-Neolithic hunter-gatherer populations from northern Austroasiatic (AA) speaking group that occupied the MSEA region prior to the migrations of East and South Asian populations ^2,16-19^.

This pattern aligns with the demographic history of MSEA, which is characterized by a AA-speaking basal layer, followed by subsequent migrations of Sino-Tibetan and Kra-Dai speakers from Southern China into northern MSEA ^18^. Furthermore, South Asian-related ancestry, associated with Indo-European speakers, introduced additional admixture signals, particularly among southern AA groups such as the Aslian-speaking Maniq. Together, these admixture events reflect an increasing genomic complexity shaped by migration routes, language shift, and cultural transitions. Therefore, self-reported ethnicity or linguistic affiliation alone is insufficient to capture the fine-scale genetic heterogeneity observed in modern Thai and MSEA populations.

By integrating multiple analytical frameworks, namely multiple linear regression (MLR) of cell proportion, differential abundance (DA) of cell states, and cell-type-specific differential expression (DE), our study identified robust associations between specific ancestry components and innate immune cell profiles in the Thai and other AIDA cohorts. More specifically within the Thai AIDA cohort, we observed significant correlations between the Blue ancestry component and gene expression variation in CD14^+^ monocytes, including up-regulation of several pro-inflammatory and interferon response genes (e.g., *IFITM2, C5AR1, MS4A7, LILRB1*). This Blue ancestry-specific expression profiles, together with high burden of tropical pathogens endemic to the region ^20,21^, suggest that MSEA-indigenous lineages might carry specialized evolutionary adaptations in innate immunity, potentially conferring a fitness advantage against regional infectious pressures ^22,23^. Our hypothesis is further supported by the known functional profiles of genes positively correlated with increasing Blue ancestry, most notably *LILRB1*, a known modulator of inflammatory responses and cytotoxicity of myeloid cell maturation of transitional monocytes against human cytomegalovirus and *Plasmodium falciparum* malaria ^24,25^, and a MHC gene (i.e., *HLA-DQB1*). This HLA gene has been identified as a locus under strong positive selection in MSEA populations and is associated with a broad spectrum of immune-related traits and disease ^4^.

Beyond the association between genetic ancestry and innate immune cell profiles, our covariate-corrected MLR and DA analyses successfully recapitulated the previously reported Thai-specific sex bias in CD16^+^ NK cell proportions ^13^. In general, sex differences in immune response in populations are shaped by X-linked genes and gonadal steroids ^26^. Our study uncovered additional unique sex-specific characteristics in the Thai population compared to other AIDA populations from both innate (i.e., CD16^+^ monocytes and CD16^+^ NK cells) and adaptive immune cells (i.e., naive CD8^+^ T cells). Specifically, XX Thai individuals exhibited greater interferon gamma responses in CD16^+^ monocytes (e.g., *SIGLEC1*.) and more highly activated states of CD16^+^ NK (e.g., up-regulation of *TNFRSF4, CD69, JUN*), whereas Thai XY group exhibited prominent gene signature of gamma-interferon-induced programmed cell death (e.g., *DAPK1*) ^27^, which was associated with a relative depletion of the TNF/NF-κB signaling pathway. In addition, we identified another Thai-specific gender-biased profile in naive CD8^+^ T cells, which were more abundant in XY individuals. Along this line, we observed up-regulation of *KLRC3* in CD8^+^ T cells of the Thai XY individuals, potentially reflecting a suppression of self-reactivity by maintaining immune homeostasis through peripheral tolerance ^28^. These together might, at least in part, indicate a unique immune phenotype or a distinct inflammatory capacity, potentially contributing to the sex-associated disease susceptibilities such as autoimmune diseases observed within the Thai and MSEA populations. These findings are also consistent with previous reports showing higher prevalence of systemic lupus erythematosus and rheumatoid arthritis in females compared to males in Thailand ^29,30^.

We next investigated whether the unique association between Blue ancestry and innate immune profiles observed in the Thai cohort persisted in other populations. The SG Malay and SG Chinese cohorts are the most ancestrally comparable to the Thai population, primarily comprising Blue and Red components. However, the ancestral association was not detectable when these cohorts were analyzed separately. This is likely because the Thai cohort uniquely features a continuous admixture gradient and broad spectrum of ancestry proportions, providing the statistical power to resolve these ancestry-driven immune patterns. Consistent with the three distinct ethnic groups in Singapore ^31^, the observed genetic components of SG populations are more polarized, with SG Malay being primarily Blue, SG Chinese being primarily Red, and SG Indian being primarily Orange - thereby lacking the intermediate mixing required to resolve this interaction. By pooling the three SG cohorts, we were able to identify a consistent association between increasing Blue ancestry and transcriptional variation in CD14^+^ monocytes across with the AIDA Thai cohort. We observed a consistent HLA gene (*HLA-DQB1*) but distinct up-regulation of inflammatory and interferon-induced genes of CD14^+^ monocytes (e.g., *TNF, MARCO, IFITM3*). Notably, the *MARCO* facilitates macrophage responses against intracellular pathogens and mediates crosstalk between innate and adaptive immunity. Genetic variation at this locus has been associated with immune diversity in Island Southeast Asian (ISEA) populations and may be influenced by historical selective pressures from *Mycobacterium tuberculosis* (TB) exposure ^3,32^. Beyond monocytes, we also observed several ancestry-associated genes of innate-like T cells (e.g., MAIT, γδ T) in the combined SG cohorts. The up-regulation of *KLRD1* in T cells have been reported in the lung of TB-infected mice and macaques ^33-35^. Given the high burden of TB in the southern Thailand, Malaysia, and Indonesia ^20,36^, this provides further evidence on the pathogen-driven adaptation to region-specific environmental factors of populations residing at Malaysian Peninsular and ISEA, that shapes distinct patterns of immune diversity compared with modern MSEA populations.

While this study provides a high-resolution map of immune diversity, several limitations remain. First, all AIDA Thailand volunteers were recruited within the Bangkok metropolitan area. While these participants represent a broad cross-section of migrants from all regions of Thailand, future studies should include rural and isolated ethnic groups to fully represent the regional diversity of MSEA countries. Second, our models could not account for all lifestyle factors, such as smoking, BMI, or historical vaccination and pandemic exposure, all of which are known to shape long-term immune landscapes and may introduce residual confounding. Finally, despite rigorous monitoring and the mitigation of batch effects across AIDA study sites, some degree of confounding may persist. To complement the comprehensive quality control process described in the AIDA marker publication ^13^, we implemented additional regression models and conducted MLR, DA and DEG analyses for each cohort independently. This stratified approach ensures that the identified ancestry-immune associations are internally consistent and not driven by site-specific technical variance. Nevertheless, further functional studies and longitudinal cross-sectional validation will be required to definitively verify these biological insights. Together, this study establishes the first representative immunological baseline for present-day Thai and MSEA populations, providing a vital framework for personalizing medical interventions in these historically underrepresented groups. This reference atlas also serves as a cornerstone for the next generation of precision medicine, such as enabling population-specific vaccine design and refining treatment predictions.

## Materials and Methods

### Study participants and sample collection

Representative “healthy” donors are a part of the AIDA Phase 1 Data Freeze v2 ^13^. In brief, a total of 619 participants were from Japan (n = 149), Singapore (n = 216), South Korea (n = 165), India (n = 30), and Thailand (n = 59). Additional PBMC samples from six different European individuals were included in each batch (n = 20) across the study sites as controls for batch effects. All participants provided written informed consent for blood sample collection, donor metadata, and clinical data, as described in the “AIDA marker paper” ^13^.

For the Thai AIDA cohort, the study was approved by the Institutional Review Boards (IRBs) of the Faculty of Medicine Siriraj Hospital, Mahidol University: IRB 725/2563 (IRB3). Study participants were recruited in Bangkok, Thailand. Eligibility was defined by Thai nationality, with no exclusion criteria regarding specific ethnic background or place of birth. The protocol for Peripheral Blood Mononuclear Cell (PBMC) isolation of all donors was consistent across all the AIDA cohorts and study sites (https://www.protocols.io/view/pbmcs-isolation-from-cpt-tube-b8r9rv96).

Inclusion criteria of sample selection for the Thai and AIDA participants required ability to provide informed consent and the absence of chronic metabolic or other diseases (e.g., diabetes (HbA1c < 6%), dementia, hypertension, asthma, auto-immune disease). Participants who had received vaccination or experienced infection within eight weeks prior to the blood collection were excluded. Other inclusion and exclusion criteria, including cell viability and quality, were described in the AIDA marker publication. Based on these criteria, the 59 PBMC samples from healthy Thai volunteers were selected as representative in the AIDA cohorts.

### Data generation and imputation of genotyping data

The protocol for genomic DNA isolation has also been described previously in the AIDA marker publication^13^. In brief, genomic DNA was extracted from PBMCs using the QIAamp DNA Mini Kit (Qiagen, cat. no. 51306). Genotyping was performed using the Illumina GSAv3.0 array comprising 654,027 fixed markers (Infinium Global Screening Array-24 Kit, cat. no. 20030770), which were aligned against the GRCh38 human genome reference. Sample and variant-level quality control of genotyping data across all AIDA samples was performed using PLINK, as described in the AIDA marker publication ^13^. After quality control, 495,589 autosomal and 21,176 X chromosomal variants among 637 samples were imputed using the Michigan Imputation Server with 2,504 human genomes from the 1000 genome project (GRCh38). For ancestry estimation, we utilized autosomal variants derived from both direct genotyping and high-density imputed data, unless indicated otherwise.

### Pre-processing steps of genotyping data between AADR and AIDA

For ancestry analysis described in **Figure S27**, we obtained modern and ancient DNA data from the Allen Ancient DNA Resource (AADR) ^15^ in the PLINK format. Genomic coordinates in Genome Reference Consortium Human Build 37 (GRCh37) were converted to Genome Reference Consortium Human Build 38 (GRCh38) using the LiftOver as part of CrossMap ^37^. After liftover and removing invalid alternative alleles, ∼71.3 % of variants (879k of a total of 1.24 million variants among 4,058 modern people) and ∼79.1% of variants (462k of a total of 584k variants among 4,316 ancient samples) passed our QC criteria. These datasets were merged using PLINK v1.9 ^38^. to identify overlapping autosomal variants. To minimize the impact of missing data, we retained only those variants with a genotype call ratio exceeding 0.99, resulting in a high-quality subset of 350,353 autosomal variants. Next, sample-level filtering for the reference panel was conducted following AADR recommendations. We excluded individuals with call rates below 0.99 or those flagged as “Ignore” or “Outlier” in the source metadata. The related or duplicated samples (KINSHIP > 0.177) were further removed using PLINK v2.0 ^39^.

The genotyping data across all AIDA samples was obtained from ^13^. A parallel QC pipeline was applied to the AIDA samples to identify and exclude related and duplicated individuals. To mitigate potential strand-alignment errors in the imputed AIDA genotyping data, all ambiguous SNPs (A/T and C/G) were filtered out. Ancestry proportions were inferred using a high-density set of 41.9 million autosomal variants across the 607 AIDA samples, benchmarked against a reference panel of 6,380 individuals (350k variants).

The unrelated AIDA samples and the AADR reference panels were merged into a single dataset. To ensure the independence of markers for ancestry inference, we performed Linkage Disequilibrium (LD) pruning using a sliding window approach by PLINK v1.9 ^38^. SNPs were removed if they exceeded an r^2^ threshold of 0.3 within a 100-kb window (step size of 5 SNPs). This procedure yielded a final set of 113,832 LD-independent autosomal SNPs across a total of 6,987 samples of AIDA (n=607) and AADR (n=6,380) for downstream ancestry inference.

### Genetic ancestry estimation

We estimated genetic ancestry using principal component analysis (PCA) through PLINK v1.9 ^38^ and global ancestry inference by ADMIXTURE v1.3.0 ^40^. PCA was conducted using the LD-pruned variants to identify broad population clusters and to project AIDA samples onto the established genetic variation of the global reference panel.

Genetic ancestry was further quantified using ADMIXTURE. The merged dataset was subset for South, East, and Southeast Asian populations. Only ethnicities represented by at least five individuals were included, resulting in a shared set of 113,832 SNVs across 2,734 individuals. We tested a range of ancestral components from K=1 to K=12. The optimal value of K was determined by identifying the model with the minimum cross-validation (CV) error. To quantify admixture patterns, the genetic diversity of each ethnicity was calculated from the ancestral components with *shannon*.*entropy()* using the following equation.

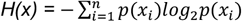

where *H(x)* is shannon entropy and *p*(*x*_*i*_) represents the mean proportion of component *x*_*i*_

### Immune cell types and immune proportional analysis among populations by single-cell transcriptomics

Immune profiles of the Thai and AIDA cohorts analysed using scRNA-seq were established and described in the AIDA marker paper ^13^. AIDA gene markers for Thirty-five immune cell types were summarized in **Figure S4**. Cell proportional analysis was calculated per individual and compared between Thai populations and other AIDA populations using multiple linear regression.

Only comparison groups with at least 50 unrelated individuals were included. Individuals with fewer than 800 cells were excluded, and only cell types with sufficient numbers of cells were included (*i*.*e*., more than seven cells per donor on median) **(Figure S28)**. Only covariate factors that have no missing data were included in the multiple linear regression models including Ancestry proportion, age, and sex. The detailed information about regression model and statistical testing were provided in the figure legends of each specific analysis.

### Differential abundance (DA) analysis and neighborhood-based differential gene expression

DA analyses were performed using MiloR v2.3.1 ^41^ to investigate the associations of genetic ancestry, age, and sex with immune cell architecture. These analyses were executed across two distinct data scales. Initial global immune profiles were derived from the established AIDA cohort datasets ^13^. Subsequently, higher-resolution DA testing was conducted by reintegrating cells using reciprocal principal-component analysis (RPCA) algorithm in Seurat v.5.2.1 ^42^ into four distinct lineage-specific objects of only individuals with more than 800 cells. These lineage-specific integrations were performed separately for myeloid innate cells (monocytes and dendritic cells), innate lymphoid and natural killer cells, T cells (double negative T, γδ T, CD4^+^ T, and CD8^+^ T cells), and B cells (B and plasma cells).

For the preprocessing step, k-nearest neighbor (KNN) graphs were constructed using a different number of k based on sample size in each study group (*i*.*e*., number of samples x 5). To ensure computational representativeness, neighborhoods were defined via random subsampling at p = 0.05 for groups exceeding 100,000 cells and p = 0.1 for smaller cohorts. The number of cells in the neighborhoods were counted.

DA testing was performed using generalized linear mixed models with spatial false discovery rate (FDR) control of the graph by Benjamini-Hochberg procedure at 0.05. We annotated immune cell types in each neighborhood using the same cell type annotations as the earlier sections. To study the relationships of genetic ancestry, age, and sex on with neighbourhood enrichment, regression models for DA were implemented using the following formulas:

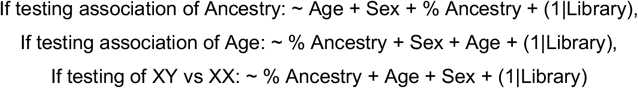

where % Ancestry (scaled from 0 to 1) and age were assigned as continuous variables. Sex coded as a categorical variable (XY and XX).

To characterize the molecular drivers of the global immune shifts, differentially expressed genes (DEGs) were identified within DA-significant neighborhoods. Expression modelling was restricted to the top 2,000 genes based on highly variable genes and performed linear models using limma v3.58.1 used in MiloR, with log-normalised counts (logcounts) as the response variable and the following model ∼ Age + Sex + % Blue ancestry as design matrix.

### Cell-type specific gene expression profiles by pseudobulk transcriptome

Differential gene expression driven by ancestral components, age, and sex for specific cell types was quantified to address how immune profiles of Thai population differs from other populations. We used the same exclusion criteria of samples and cell types as the proportional analysis above.

Genes were retained if their expression exceeded 0.5 CPM in at least 10 donors within that cell-type group. Retained counts were normalised for library size which was multiplied by normalized factor using trimmed mean of M-values (TMM) normalisation, and expression values were converted to log-CPM. Differential expression analysis was performed using a linear mixed model from Dream ^43^ within the variancePartition package v1.32.5 ^44^. The following model was applied per cell type:

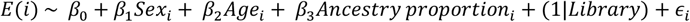

where *E*(*i*) is a log-cpm gene expression value with precision weights of gene *i* of each specific cell type. *β*_0_ is y-intercept, *β*_1_ is coefficient levels between XY and XX, *β*_2_ is the effects of age, and *β*_3_is the effects of ancestry proportion, (1|*Library*) represents random intercept for library preparation, and *ϵ*_*i*_ is the residual error. Statistical significance was assessed using the Empirical Bayes approach, and p values were adjusted using the Benjamini-Hochberg procedure. Genes with |log2FC| > 1 and adjusted P-value < 0.05 were considered as DEGs.

### Gene set enrichment analysis (GSEA)

Gene set enrichment analysis (GSEA) was performed to investigate the molecular mechanisms underlying genetic ancestry-, age-, and sex-associated immune variation. The Hallmark gene set collection for *Homo sapiens* was retrieved from the Molecular Signatures Database (MSigDB, version 25.1.1) ^45^ as the reference pathway database. Genes were ranked by the signed ranking metric: sign(log2FC) ×™log10(adjusted P-value). GSEA was conducted using the *fgsea()* function ^46^, with minimum and maximum gene set sizes of 10 and 800 genes, respectively. Normalised enrichment scores (NES) are reported with Benjamini–Hochberg-adjusted p-values. The analysis was applied in two distinct contexts: neighborhood-based enrichment utilized the ranked 2,000 highly variable genes of DA-significant neighbourhoods from the MiloR DA analysis to identify pathways associated with Blue ancestry, while cell-type-specific enrichment utilized full ranked gene lists from the pseudobulk analysis across individual of specific cell type to resolve pathways associated with ancestry, age, and sex immune lineages.

## Data, code, and materials availability

All original genotyping data and processed single-cell RNA-seq data with metadata used in this study are available via the previously published dataset (Kock et al., 2025). The AIDA Data Freeze v2 gene-cell matrix and metadata are available via CZ CELLxGENE (https://cellxgene.cziscience.com/collections/ced320a1-29f3-47c1-a735-513c7084d508) and the Cell Annotation Platform (https://celltype.info/project/336/dataset/591). The open-access dataset is available via the HCA Data Portal (https://data.humancellatlas.org/explore/projects/f0f89c14-7460-4bab-9d42-22228a91f185). Managed access datasets are available via https://explore.data.humancellatlas.org/projects/35d5b057-3daf-4ccd-8112-196194598893 and https://explore.data.humancellatlas.org/projects/76bc0e97-8cae-43d4-a647-477a13be47f9. The raw sequencing data of a Thai dataset is available from the lead contact upon request. All original code has been deposited at Github is publicly available at https://github.com/vclabsysbio/AIDAThailand_PhaseI as of the date of publication.

## Author contributions

Conceptualization: VC, PM, BS, BSr, PC

Methodology: VC, PM, BS, MP, SN, DJ, NT, BSr, PC, Asian Immune Diversity Atlas Network Investigation: VC, PM, BS, MP, SN, BSr, PC

Visualization: BSr

Funding acquisition: VC, PM, BS, MP, Asian Immune Diversity Atlas Network

Project administration: VC, PM, BS, MP

Supervision: VC, PM, BS, MP

Writing – original draft: VC, BSr

Writing – review & editing: All authors

Asian Immune Diversity Atlas Network: Varodom Charoensawan, Chung-Chau Hon, Partha P. Majumder, Ponpan Matangkasombut, Woong-Yang Park, Shyam Prabhakar, Jay W. Shin, Piero Carninci, John C. Chambers, Marie Loh, Manop Pithukpakorn, Bhoom Suktitipat, Kazuhiko Yamamoto, Deepa Rajagopalan, Nirmala Arul Rayan, Shvetha Sankaran, Juthamard Chantaraamporn, Ankita Chatterjee, Supratim Ghosh, Kyung Yeon Han, Damita Jevapatarakul, Sarintip Nguantad, Sumanta Sarkar, Narita Thungsatianpun, Mai Abe, Seiko Furukawa, Gyo Inoue, Keiko Myouzen, Jin-Mi Oh, Akari Suzuki, Yoshinari Ando, Miki Kojima, Tsukasa Kouno, Jinyeong Lim, Arindam Maitra, Le Min Tan, Prasanna Nori Venkatesh, Murim Choi, Jong-Eun Park, Eliora Violain Buyamin, Kian Hong Kock, Quy Xiao Xuan Lin, Jonathan Moody, Radhika Sonthalia, Kazuyoshi Ishigaki, Masahiro Nakano, Yukinori Okada & Yoshihiko Tomofuji.

## Acknowledgments

We thank all members of the Asian Immune Diversity Atlas (AIDA) network for their valuable comments and suggestions throughout the development of this manuscript. We are also grateful to Irene Gallego Romero and her team, for their insightful discussions and feedback throughout the submission process. We extend our sincere gratitude to all the volunteers and clinical staff whose participation and dedication made this study possible. The authors acknowledge the use of generative AI tools (ChatGPT and Google Gemini), during the preparation of this manuscript; these tools were utilized solely for grammar checks and to improve readability of the text. The authors take full responsibility for the final contents and scientific integrity of this work.

