## Supplementary Figure 1-28 for "Cellular Complexity and Systemic Immune Profiles across Ancestral Diversity in Thailand and Mainland Southeast Asia"

### Supplemental information

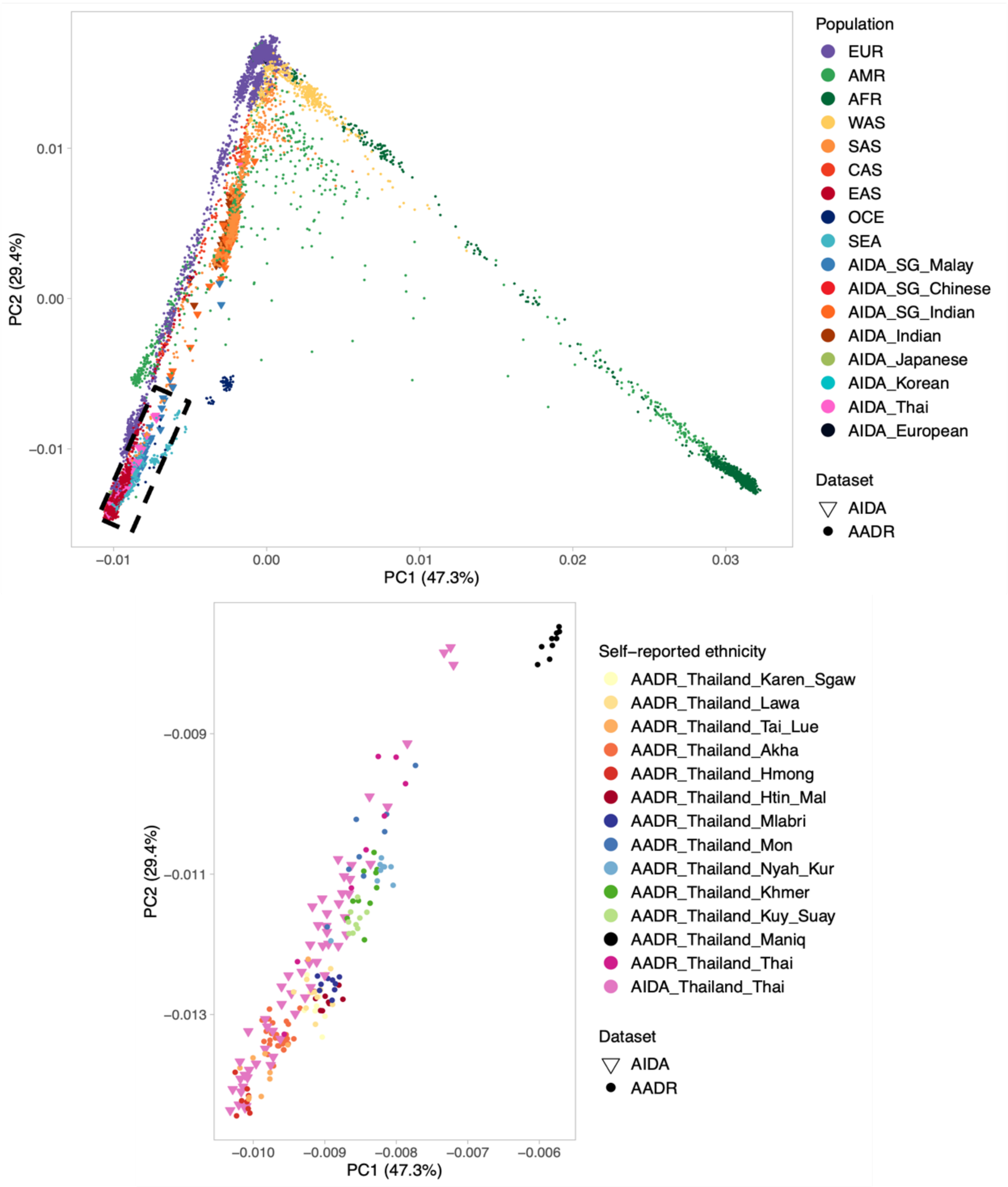


**Figure S1. Principal Component Analysis (PCA) analysis of unrelated samples across AIDA and global populations. (Top)** Genetic structure of AIDA samples and the global populations from the Allen Ancient DNA Resource (AADR). Populations from EUR, Europe; AMR, Americas; AFR, Africa; WAS, Western Asia; SAS, South Asia; CAS, Central Asia; EAS, Eastern Asia; OCE, Oceania; SEA, South-eastern Asia. **(Bottom)** PCA restricted to the majority group of 58 present-day Thai populations in AIDA and Thai reference samples. The PCA was performed using 113,832 autosomal SNPs after imputation and quality controls.

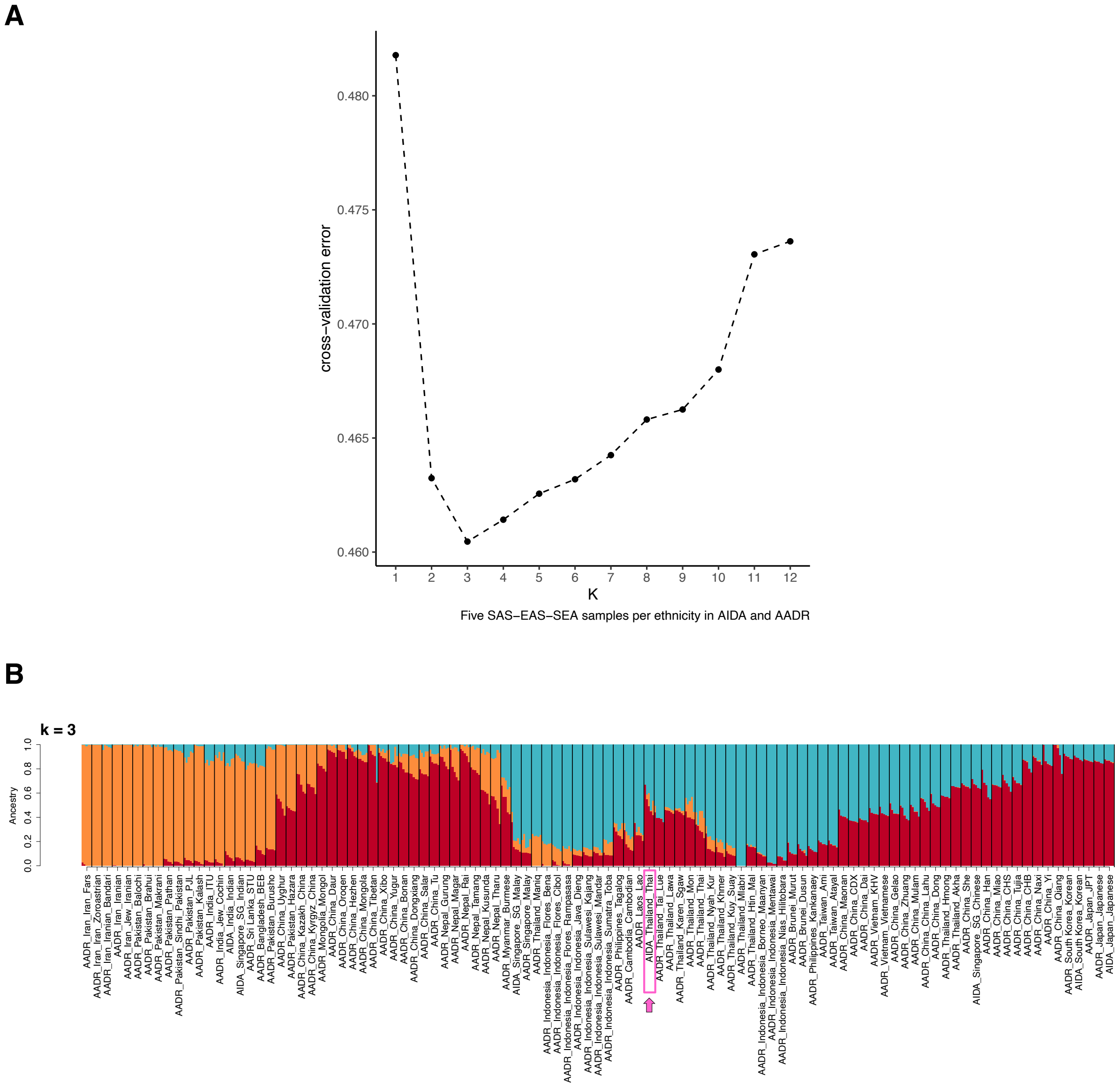


**Figure S2. ADMIXTURE analysis of individuals across the AIDA and the AADR with equal sample size per ethnicity**. **(A)** Cross-validation error estimates for k = 1 to k = 12. **(B)** Ancestral component at k = 3 per individual based on five unrelated samples per ethnicity. Thai individuals in the AIDA cohort are highlighted in pink. The analyses were performed using 113,832 autosomal SNPs after imputation and quality controls of five unrelated samples per ethnicity.

**A**

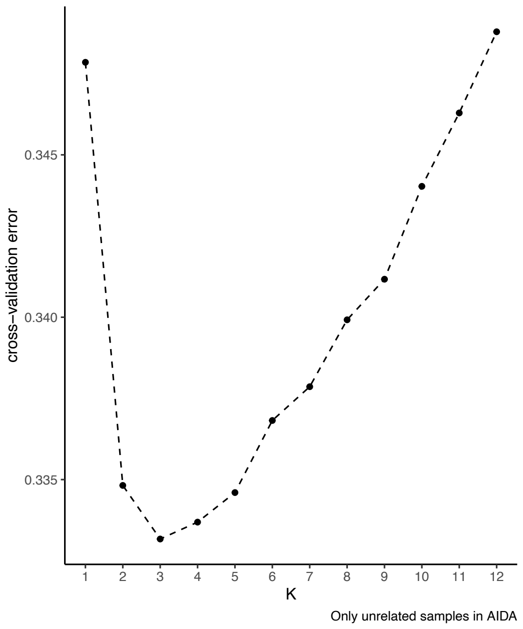


**B**

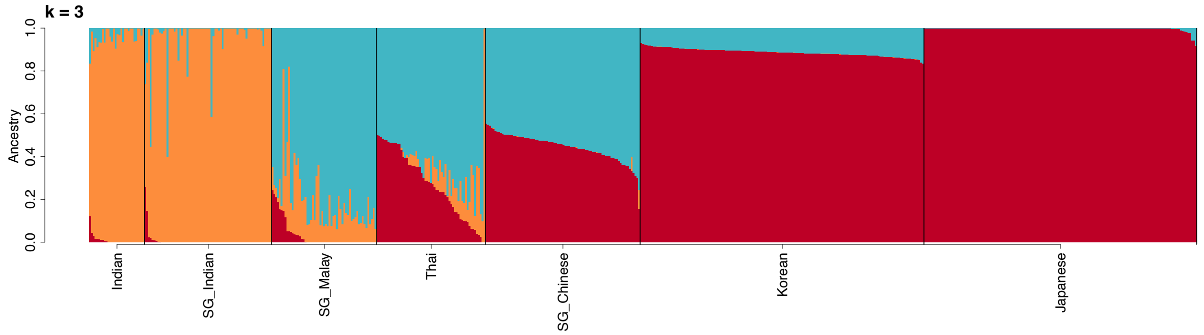


**Figure S3. Estimation of ancestral components among 601 unrelated individuals from the AIDA populations. (A)** Cross-validation error estimates from k = 1 to k = 12. **(B)** Proportion of ancestral components of each AIDA individual at k = 3. ADMIXTURE analysis was performed using 495,589 autosomal SNPs from Global Screening Array data.

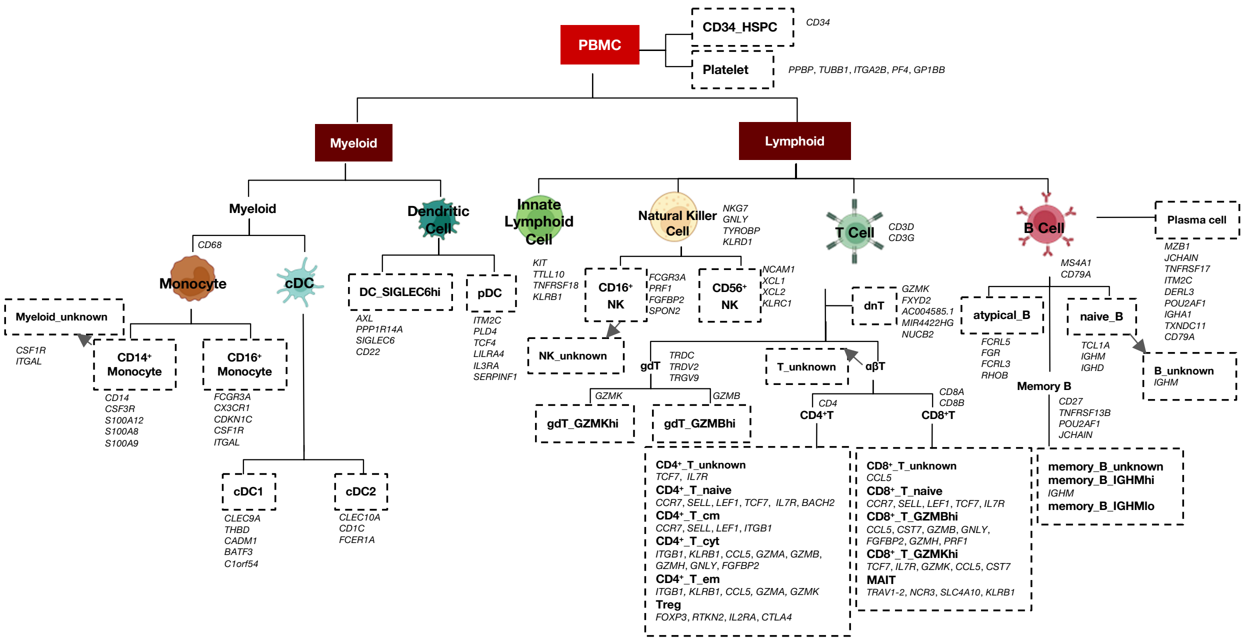


**Figure S4. Hierarchy of 35 immune cell type annotation with AIDA gene markers (see the AIDA marker publication for more details).**

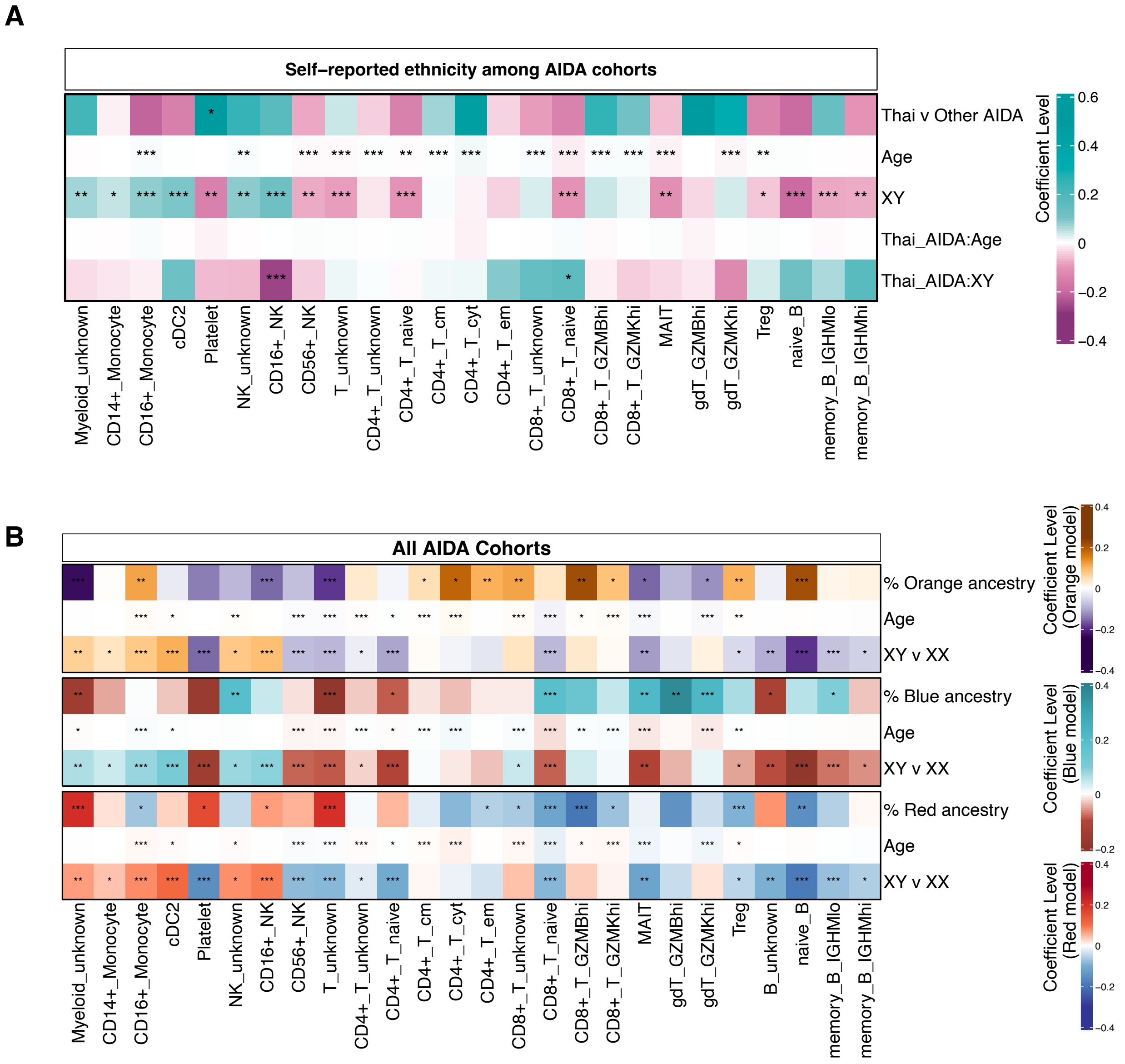


**Figure S5. Cell proportional analysis across the AIDA cohorts.** The regression models were performed using **(A)** interaction terms of Thai population compared with Other AIDA (*i.e.*, Singaporean, Korean, and Japanese) ($\log10(\text{Proportion})_{\text{cell type}}$ ~ Cohort + Age + Sex + Cohort:Age + Cohort:Sex) and **(B)** accounting for ancestral diversity $\log_{10}$(Proportion) ~ % Ancestry + Age + Sex). Statistical analyses were performed using MLR; * $p < 0.05$, ** $p \leq 0.01$, *** $p \leq 0.001$.

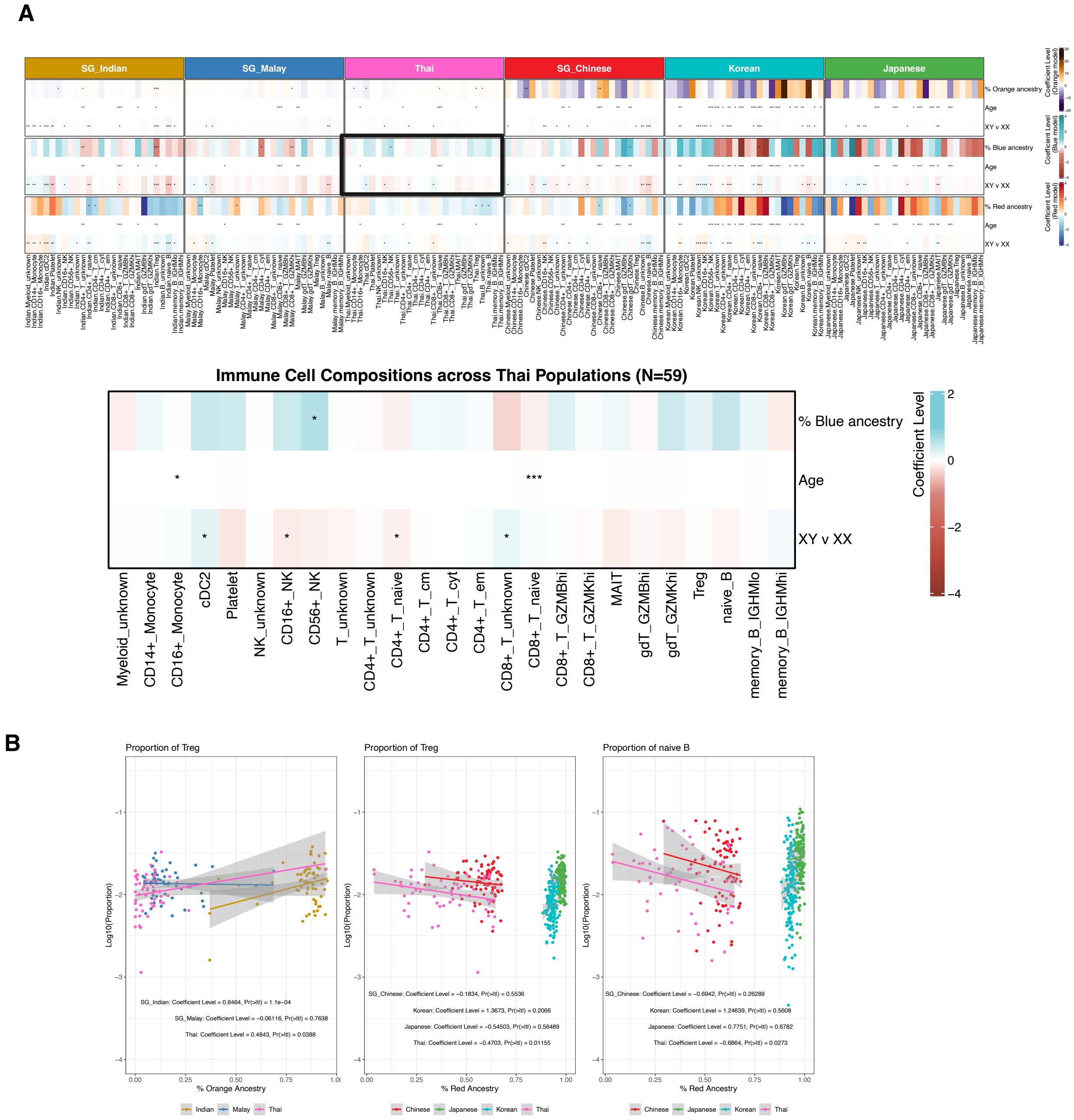


**Figure S6. Immune cell proportion analysis among the AIDA samples.** The plots presents cell proportional analysis of **(A)** each ethnicity using different ancestral components as a genetic factor ($\log_{10}$(Proportion) ~ % Ancestry + Age + Sex), with a zoomed view in the Thai cohort **(B)** The plots show correlation between the proportion of Treg and naive B cells with Red or Orange ancestral component across AIDA samples. Statistical analyses were performed using MLR ($\log_{10}$(Proportion) ~ % Ancestry + Age + Sex).

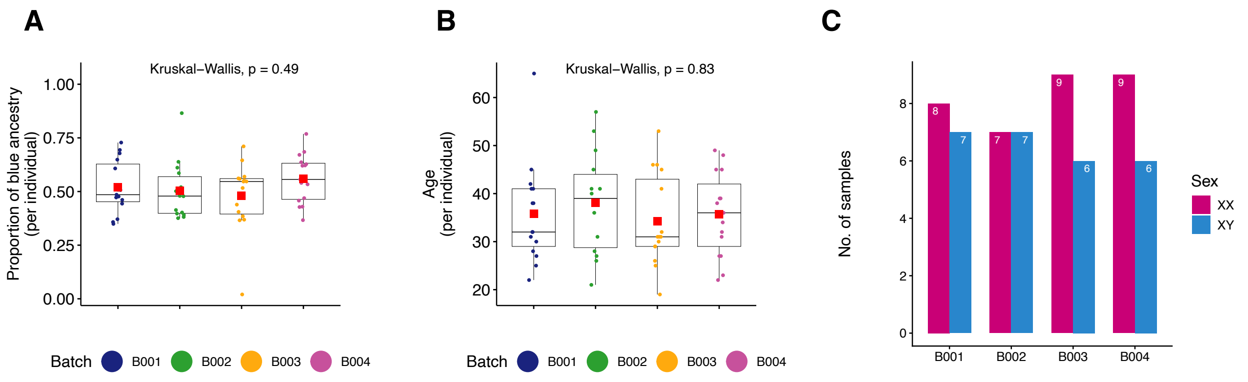


**Figure S7. Sample distributions across four batches in the Thai AIDA cohort. (A)** Distribution of proportion of Blue ancestral components of each Thai individual. **(B)** age distributions across the batches. Means of blue ancestry and age across 16 people in each batch are labeled by red square, and **(C)** number of XX and XY individuals in each batch of the AIDA Thailand cohort.

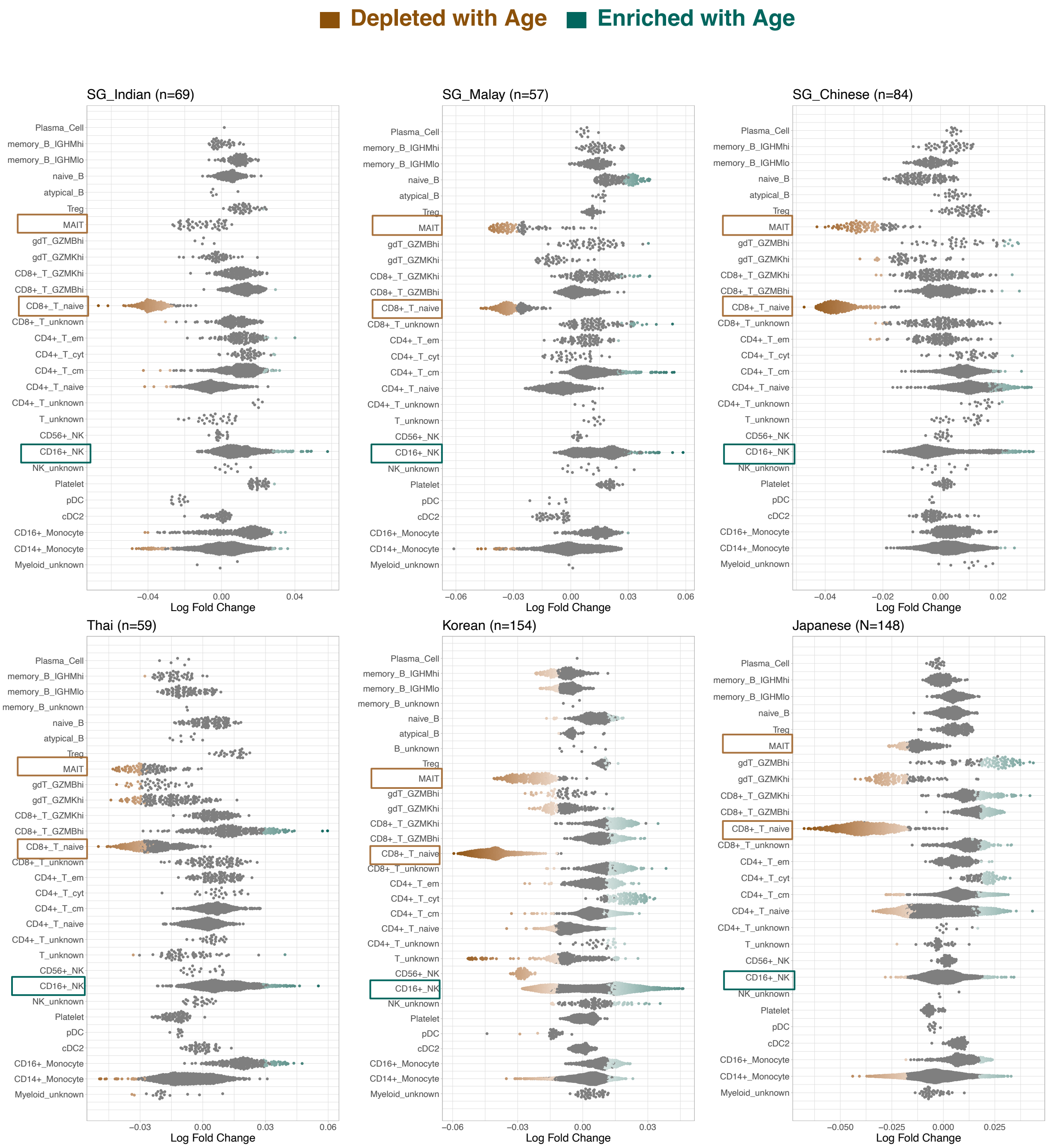


**Figure S8. Impact of age on immune cell abundances.** Beeswarm plots showing the neighborhood depletion/enrichment with age among the AIDA cohorts. The statistical testing was using GLMM per each neighborhood (Cell count$_{nhood}$ ~ Sex + % Main Ancestry + Age + (1|Library) at spatial FDR < 0.05). “Main Ancestry” used in the model were Blue for AIDA Thailand, SG Chinese and SG Malay; Red for AIDA Japan and Korea; and Orange for SG India.

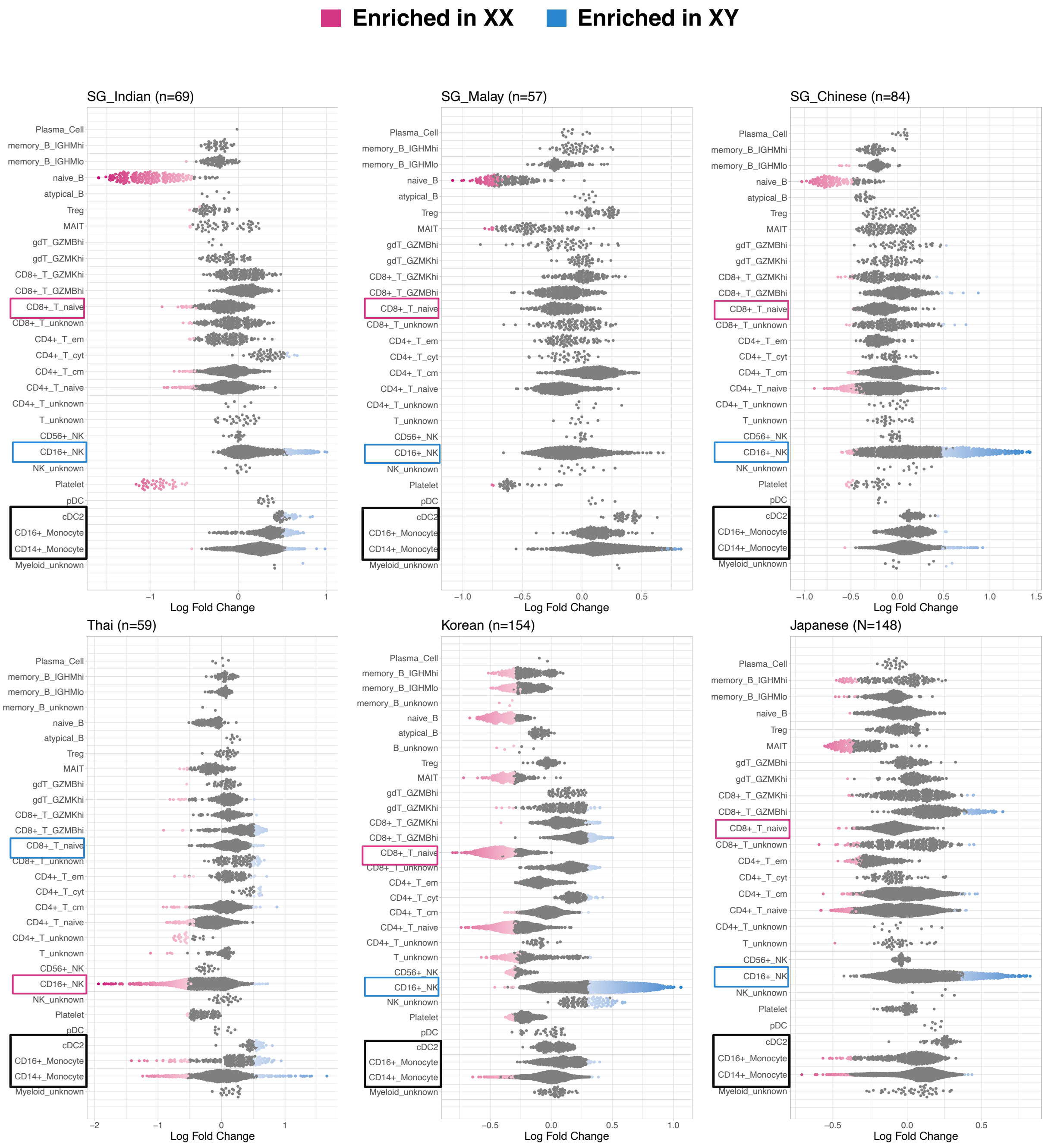


**Figure S9. Impact of sex on immune cell abundances.** Beeswarm plots showing the neighborhood depletion/enrichment with XY among the AIDA cohorts. The statistical testing was using GLMM per each neighborhood (Cell count$_{nhood}$ ~ Age + % Main Ancestry + Sex + (1|Library) at spatial FDR < 0.05). “Main Ancestry” used in the model were Blue for AIDA Thailand, SG Chinese and SG Malay; Red for AIDA Japan and Korea; and Orange for SG India.

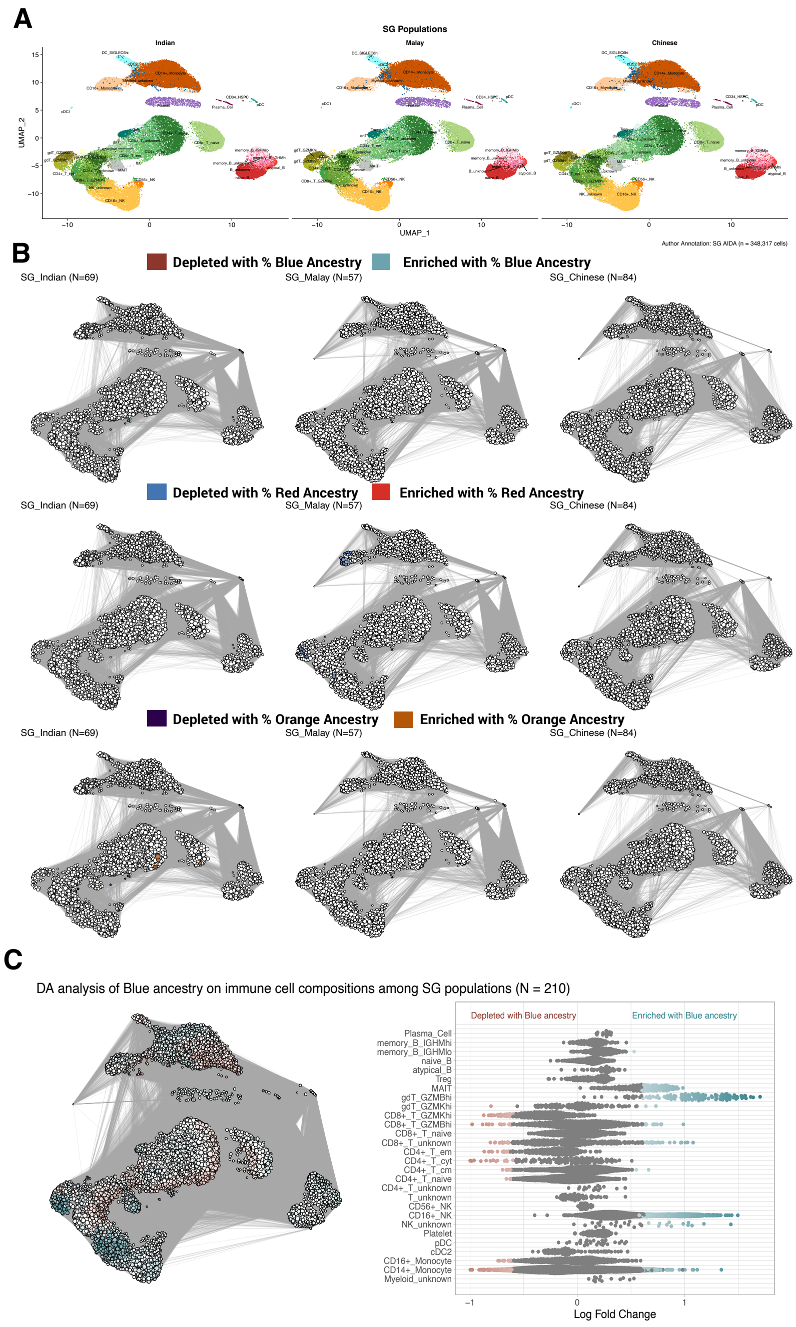


**Figure S10. Impact of ancestry diversity on differential cell abundances among the three SG AIDA cohorts. (A)** UMAP visualization of immune cell types across each SG population. **(B)** DA analysis showing association of Blue ancestry, Red ancestry, Orange ancestry among each SG population at spatial FDR < 0.05. **(C)** Overall impact of Blue ancestry on cell abundances among all Singaporeans at **(right)** spatial FDR < 1 and **(left)** the neighborhood enrichment with Blue ancestry among all Singaporean at spatial FDR < 0.05.

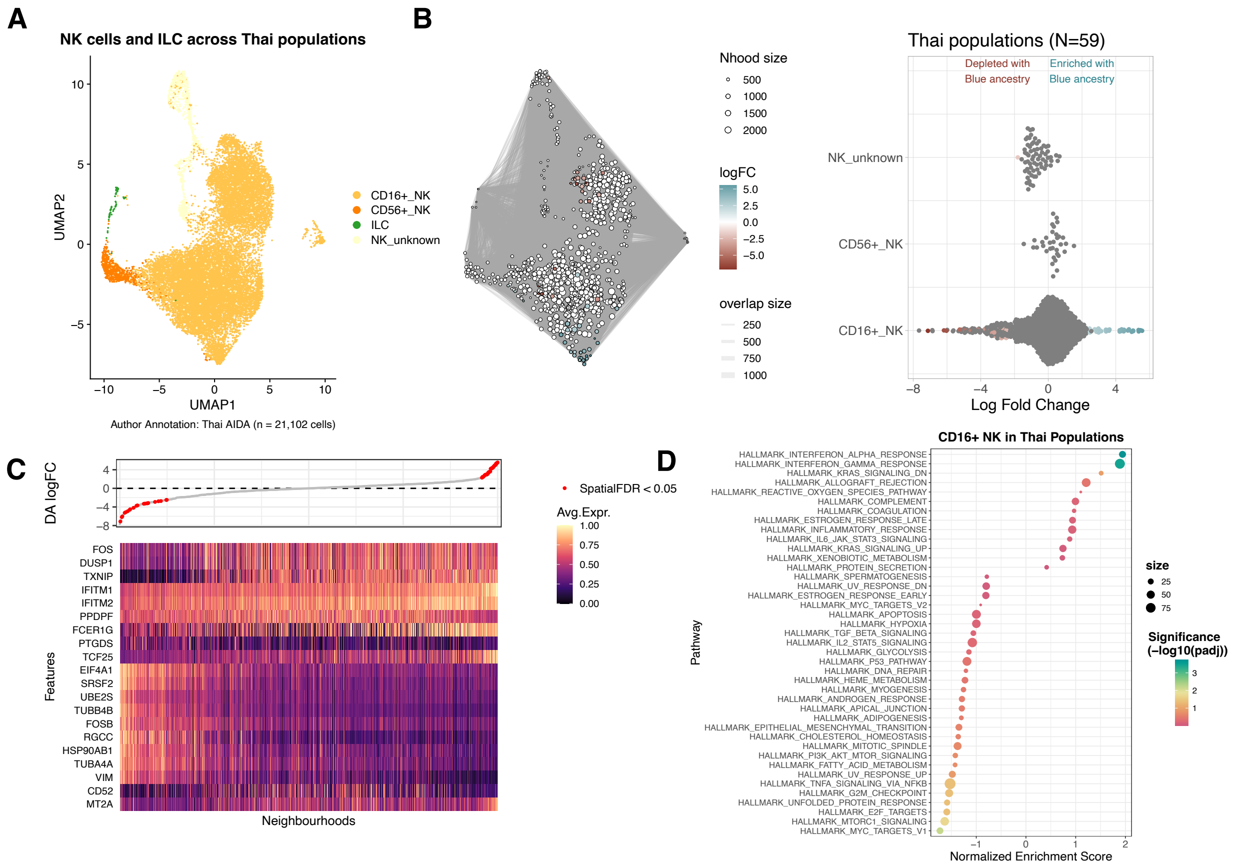


**Figure S11. The neighborhoods correlate with Blue ancestry on innate lymphoid cells and natural killer cells among the Thai AIDA population. (A)** UMAP showing cell type identification **(B)** the DA and beeswarm plots show association of Blue ancestry on differential cell abundance (DA) of neighborhoods correlated with Blue ancestry among Thai populations (GLMM: Cell count$_{nhood}$ ~ Age + Sex + % Blue ancestry + (1|Library) at spatial FDR < 0.05). **(C)** The heatmap plot shows association of Blue ancestry on differential gene expression of CD16$^{+}$ NK neighborhoods correlated with Blue ancestry (MLR: Gene expression$_{nhood}$ ~ Age + Sex + % Blue ancestry at |logFC| > 0.5 and adj.P.Val < 0.05). and **(D)** pathway enrichment by GSEA analysis across 2,000 signature genes.

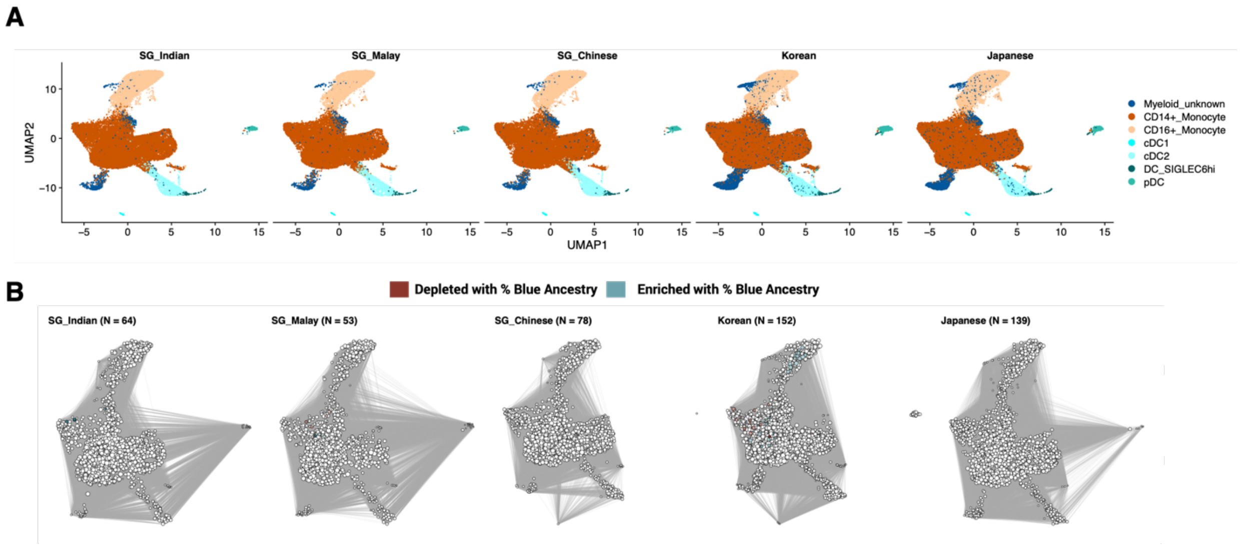


**Figure S12. Myeloid innate immune cell types among the AIDA cohorts. (A)** Uniform Manifold Approximation and Projection (UMAP) plots shows immune cell types of myeloid innate Immune cells after RPCA integration of each ethnicity. **(B)** correlation of Blue ancestry on immune cell abundances of myeloid innate immune cells among each AIDA population. The DA analysis were tested using GLMM per each neighborhood (Cell count$_{nhood}$ ~ Age + Sex + % Blue ancestry + (1|Library) at spatial FDR < 0.05).

A

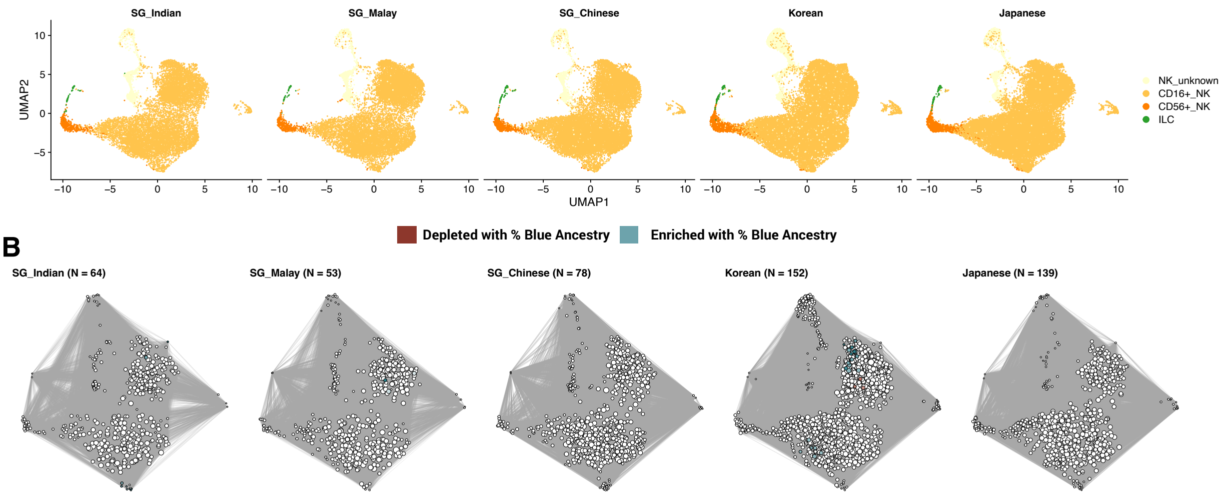


B

**Figure S13. Lymphoid innate immune cell types among the AIDA cohorts. (A)** Uniform Manifold Approximation and Projection (UMAP) plots shows immune cell types of innate lymphoid cell (ILC) and natural killer cells after RPCA integration. **(B)** correlation of Blue ancestry on immune cell abundances of NK and ILC immune cell type among each AIDA population. The DA analysis was tested using GLMM per each neighborhood (Cell $count_{nhood}$ ~ Age + Sex + % Blue ancestry + (1|Library) at spatial FDR < 0.05).

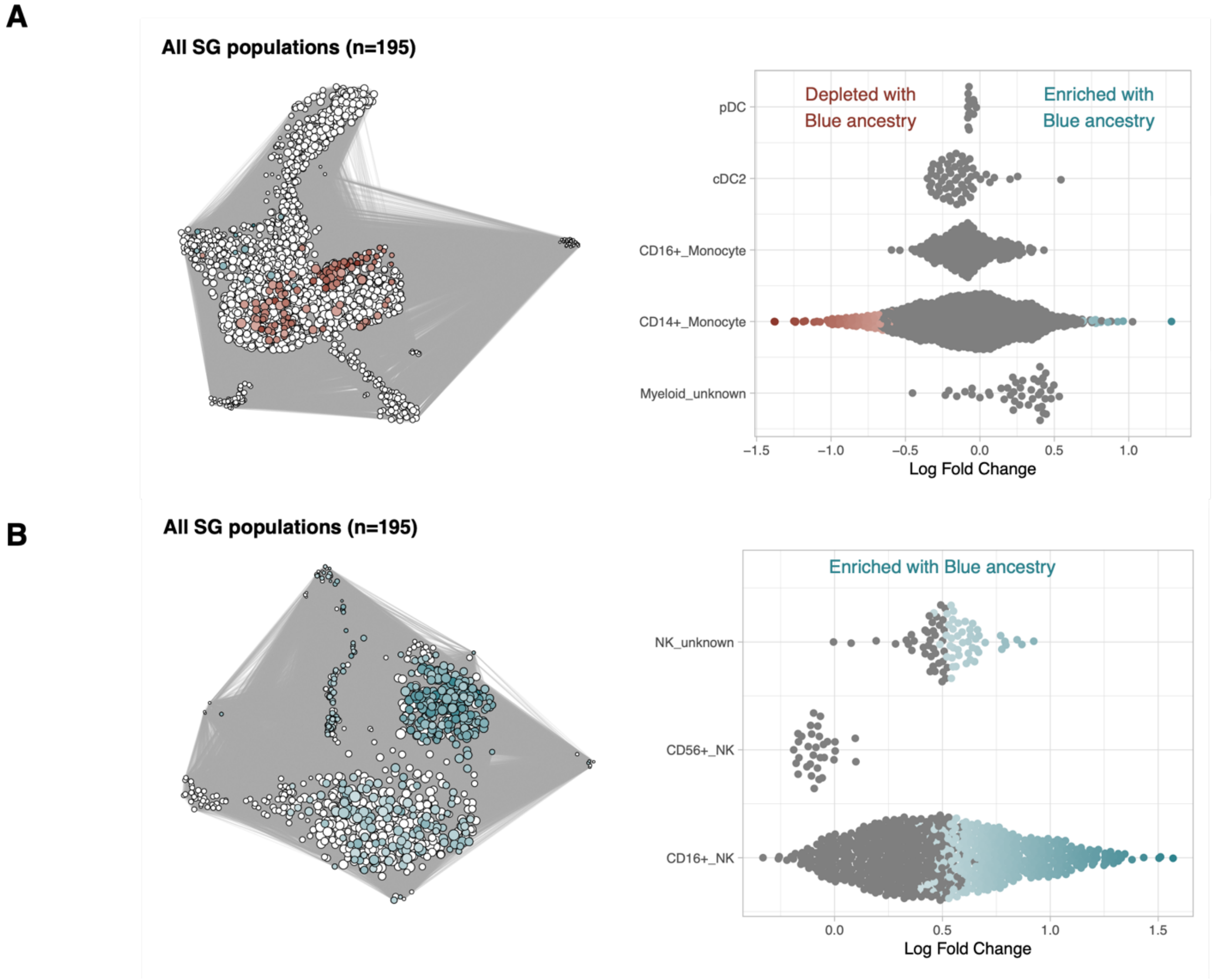


**Figure S14. Correlation of Blue ancestry on innate immune cell abundances among the SG AIDA populations**. Differential abundance analysis was performed on **(A**) Myeloid innate cells and **(B)** NK with ILC (GLMM: Cell count$_{nhood}$ ~ Age + Sex + % Blue ancestry + (1|Library) at spatial FDR < 0.05).

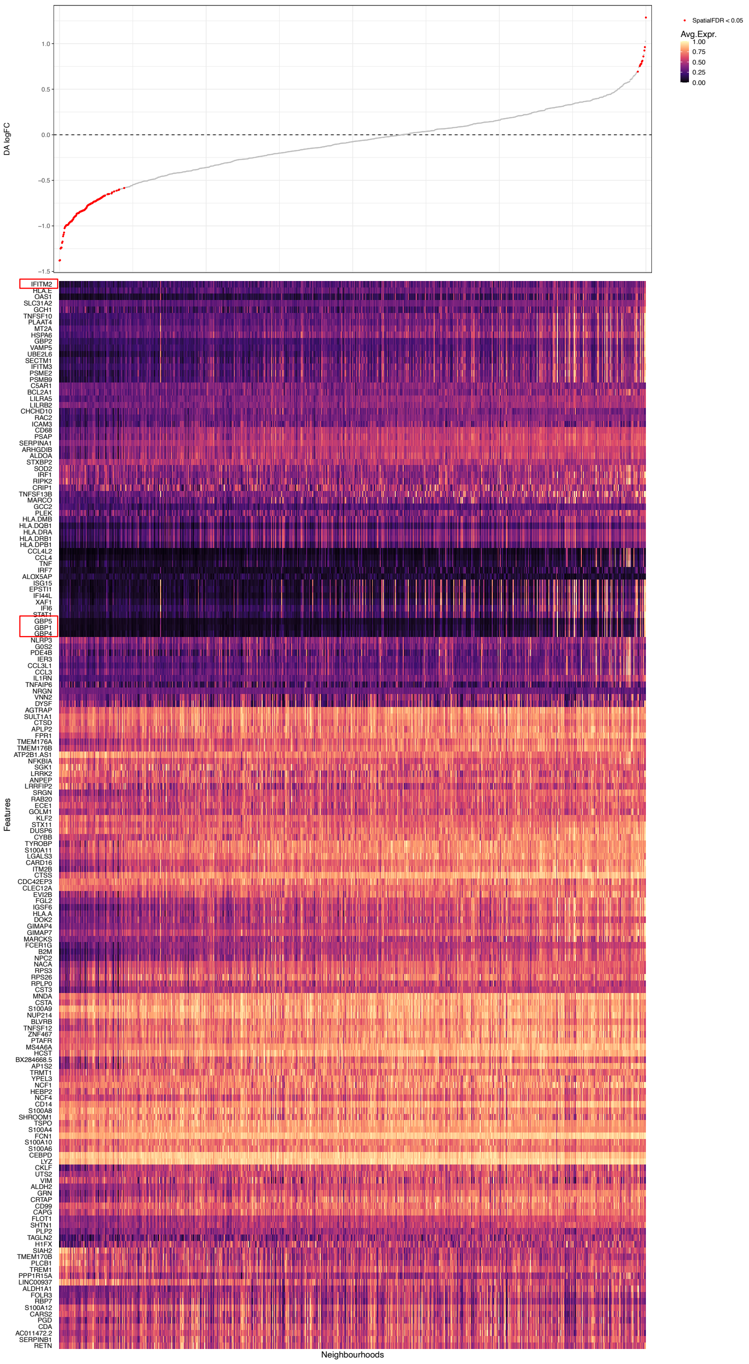


**Figure S15. The neighborhood enrichment associated with Blue ancestry and their gene expression profiles among the SG AIDA populations.** The statistical testing for DA was using GLMM per each neighborhood (Cell count$_{nhood}$ ~ Age + Sex + % Blue ancestry + (1|Library) at spatial FDR < 0.05) and The DEGs testing was using MLR per gene in each neighborhood (Gene expression$_{nhood}$ ~ Age + Sex + % Blue ancestry at logFC > 0.05 & adj.P.Val < 0.05).

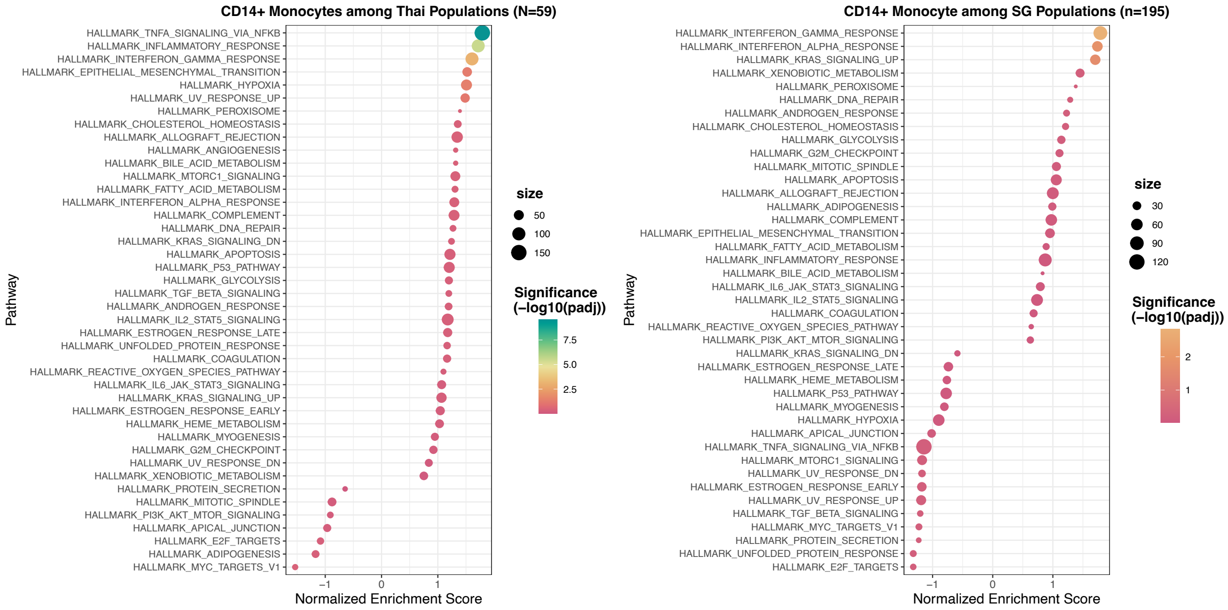


**Figure S16. Hallmark pathways of CD14⁺ monocyte neighborhoods associated with Blue ancestry by gene set enrichment analysis.** The 2,000 highly variable signature genes were performed in GSEA analysis.

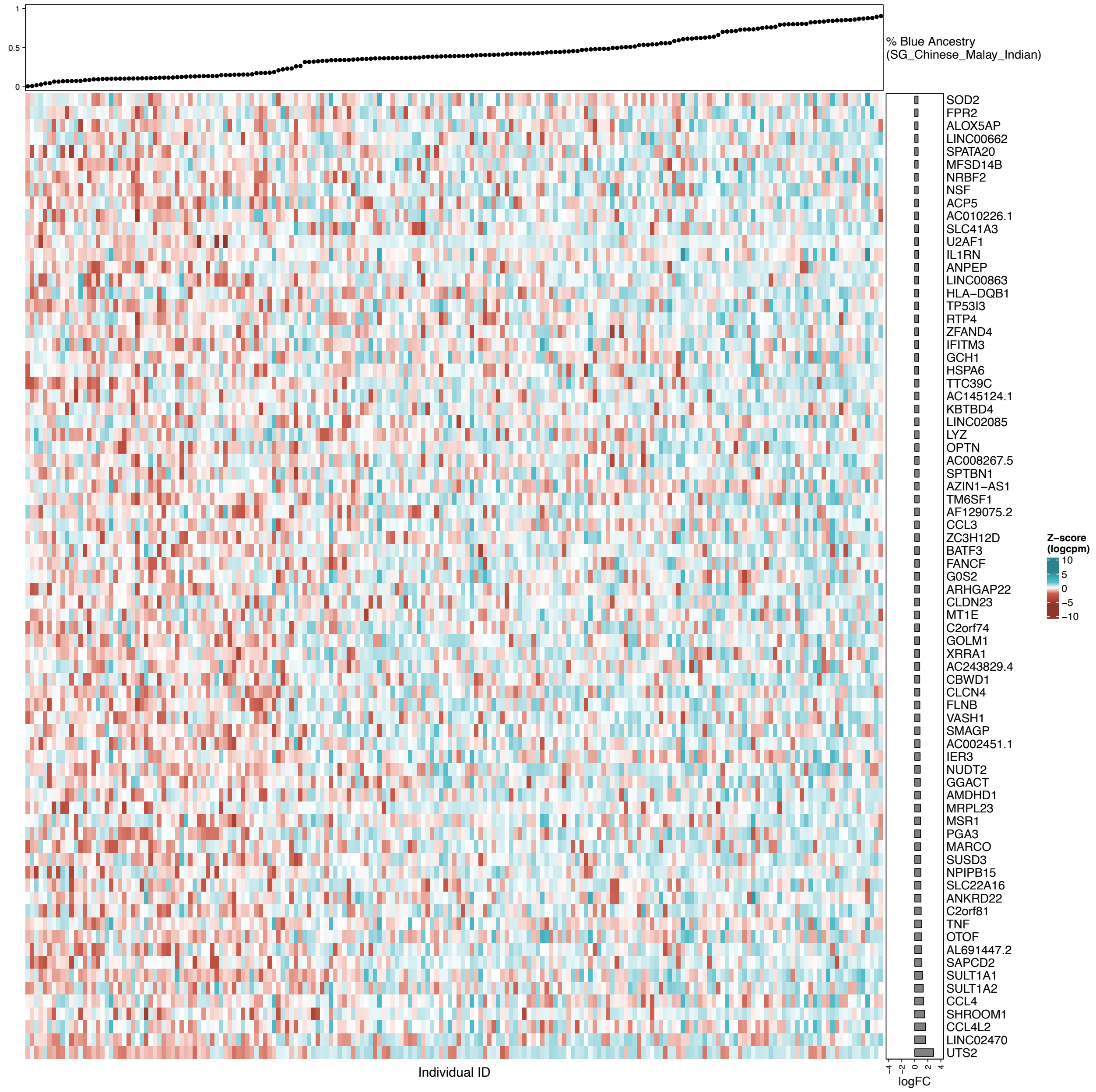


**Figure S17. Gene expression of CD14⁺ monocytes among the SG AIDA populations.** The heatmap plots showing gene expression pattern of upregulated genes associated with Blue ancestry among all Singaporeans per individual at logFC > 0.05 and adj.P.Val < 0.05.

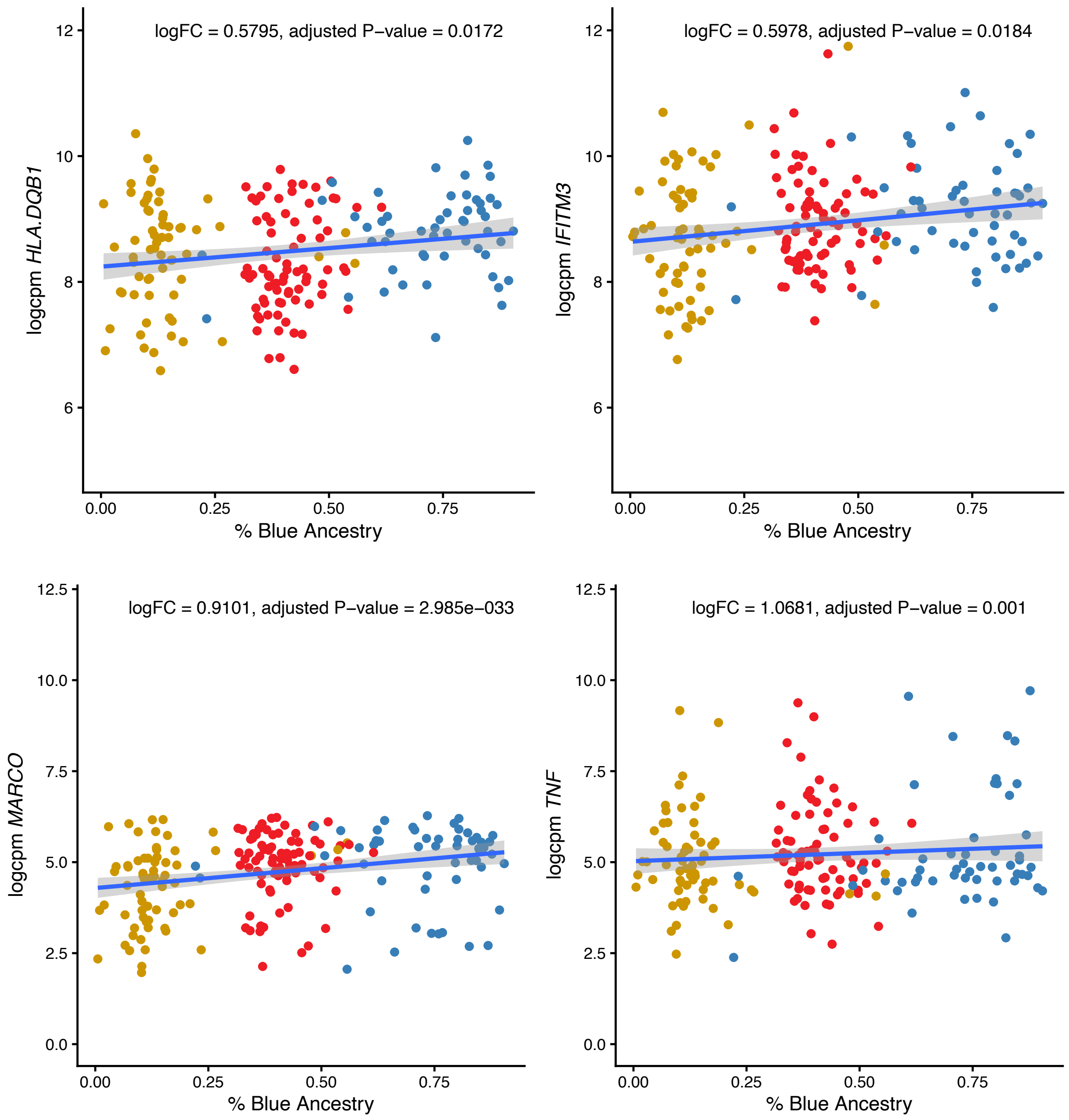


**Figure S18. The correlation of representative gene expression in CD14⁺ monocytes and Blue ancestral proportion among SG populations.**

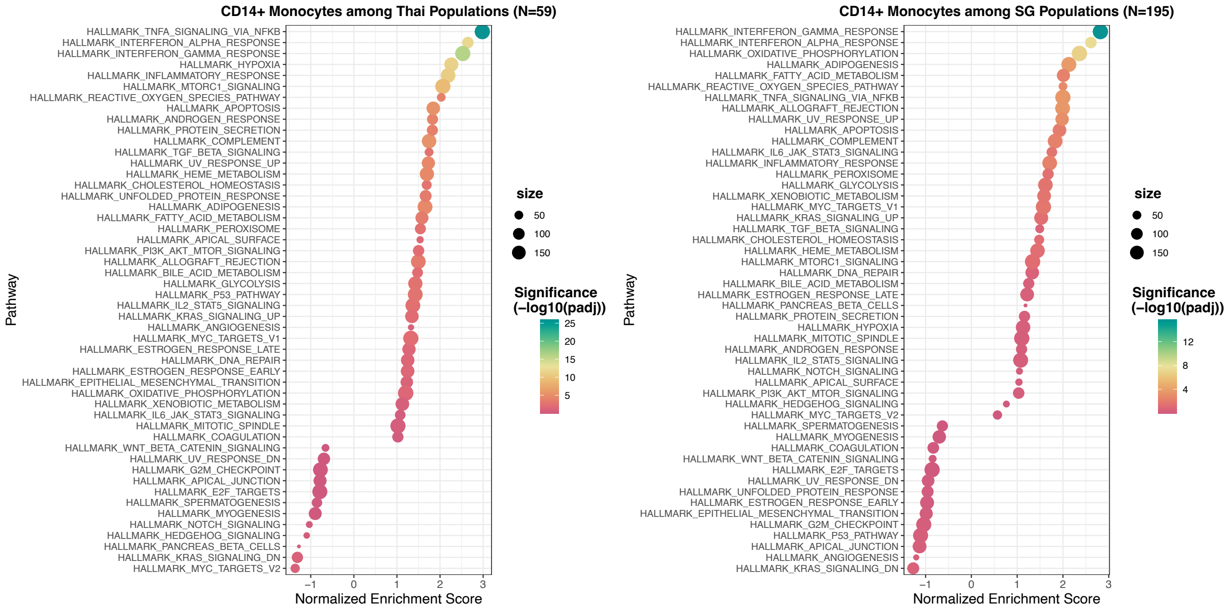


**Figure S19. GSEA analysis on pseudobulk-transcriptomes per individual of CD14⁺ monocyte associated with Blue ancestry of the Thai and SG AIDA populations.**

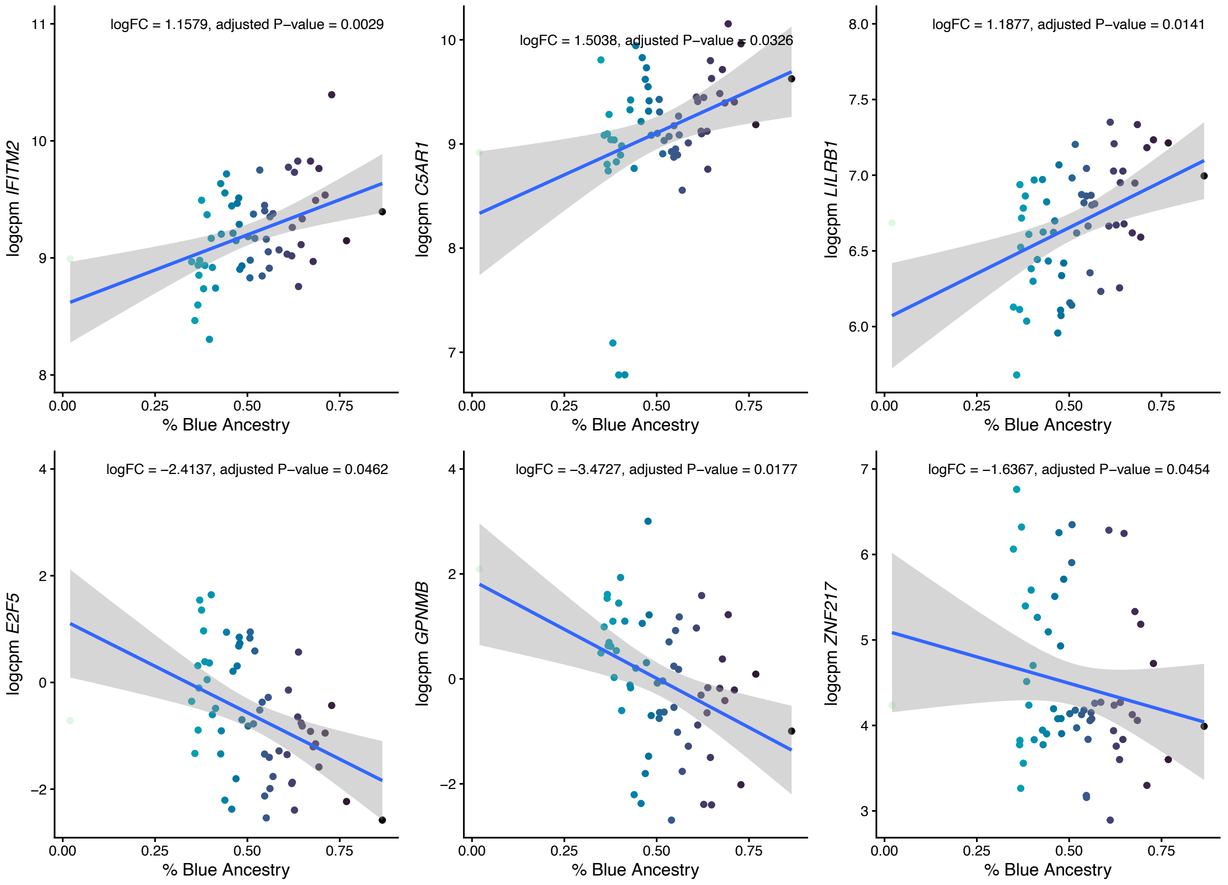


**Figure S20. Representative differentially expressed genes correlated with Blue ancestry in $CD14^+$ monocytes of the Thai AIDA population.**

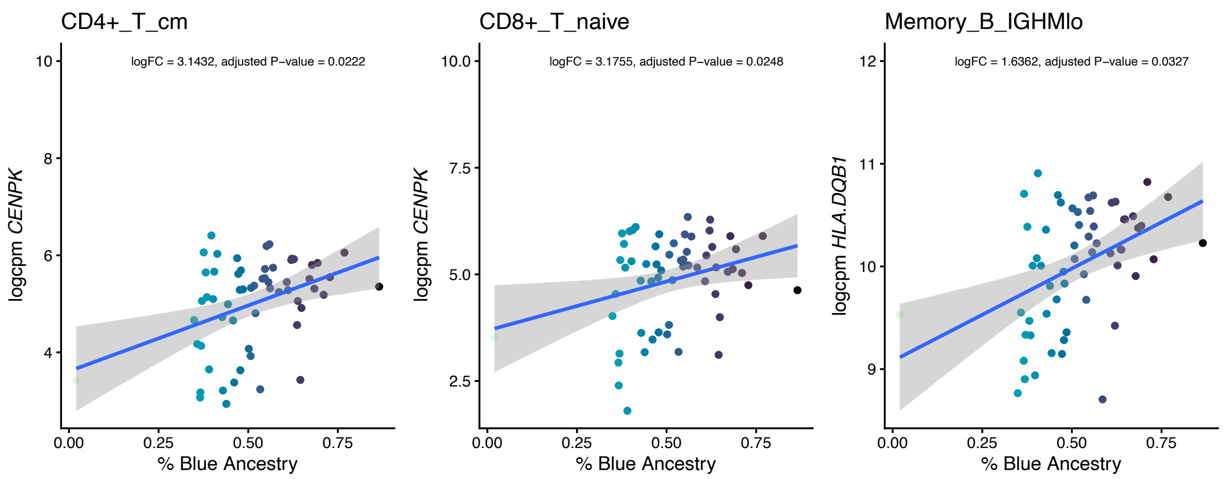


**Figure S21. Representative differentially expressed genes correlated with Blue ancestry in adaptive immune cells of the Thai AIDA population.**

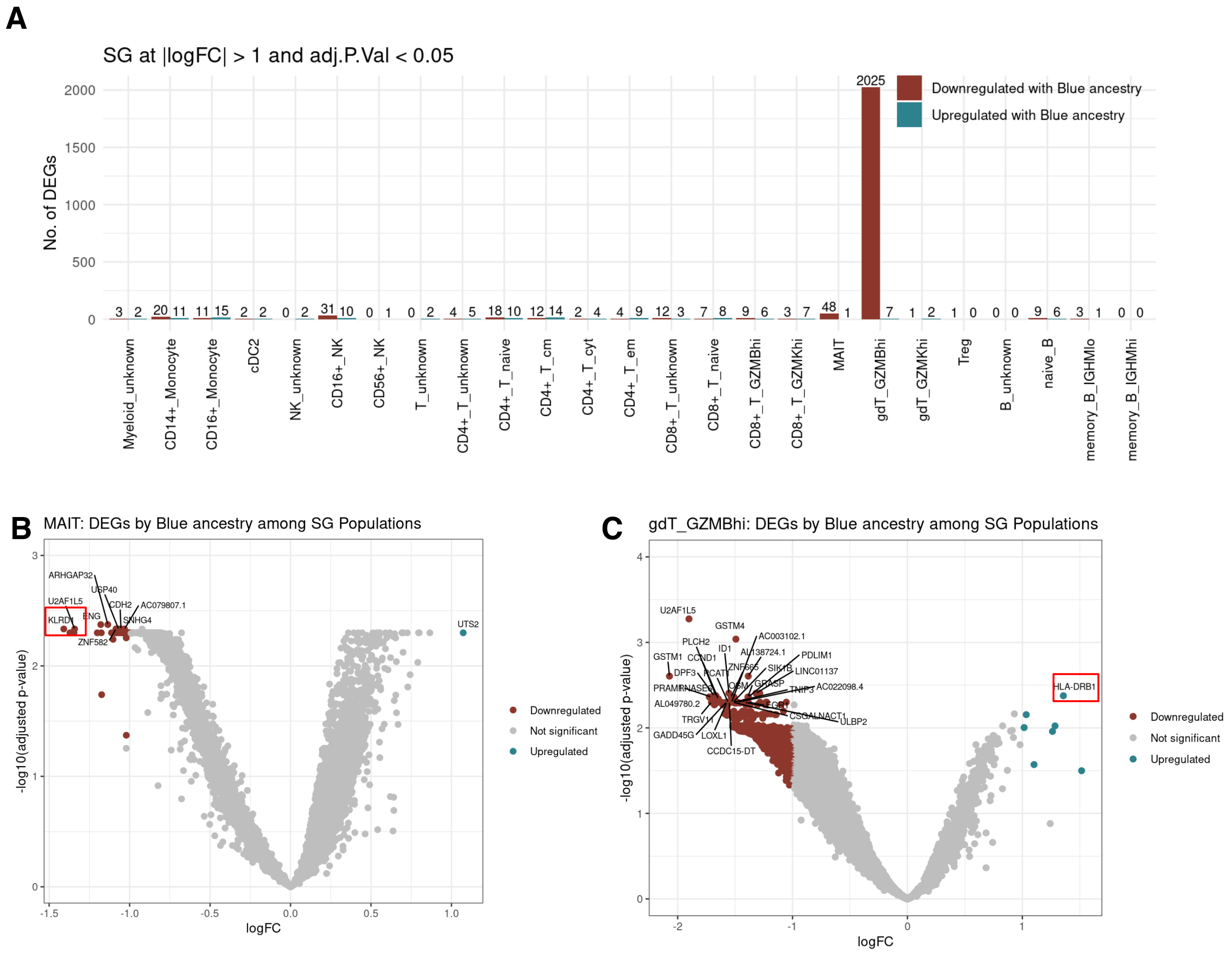


**Figure S22. Impact of Blue ancestry on expression profiles of innate and adaptive immune cells among the SG AIDA populations. (A)** The barplots represent number of DEG associated with Bue ancestry in each cell type. **(B)** DEGs of MAIT and **(C)** γδ T cells associated with Blue ancestry among combined three SG populations. The DEGs testing was using LMM per gene in each cell type of each individual (logcpm(Gene expression)$_{\text{cell type}}$ ~ Age + Sex + % Blue ancestry + (1|Library) at |logFC| > 1 and adj.P.Val < 0.05).

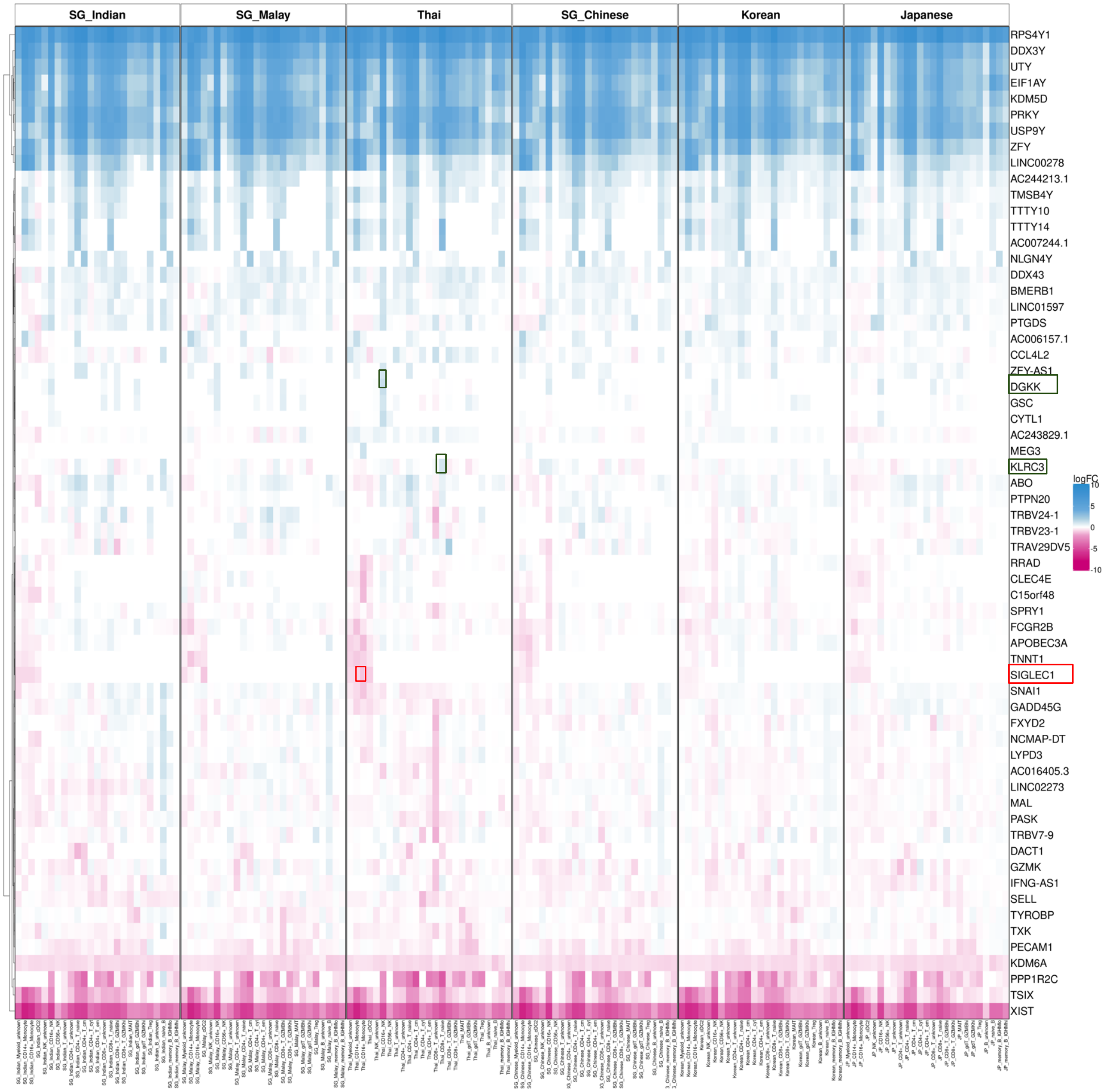


**Figure S23. Sex Differences in cell-type specific gene expression in each AIDA cohort.**

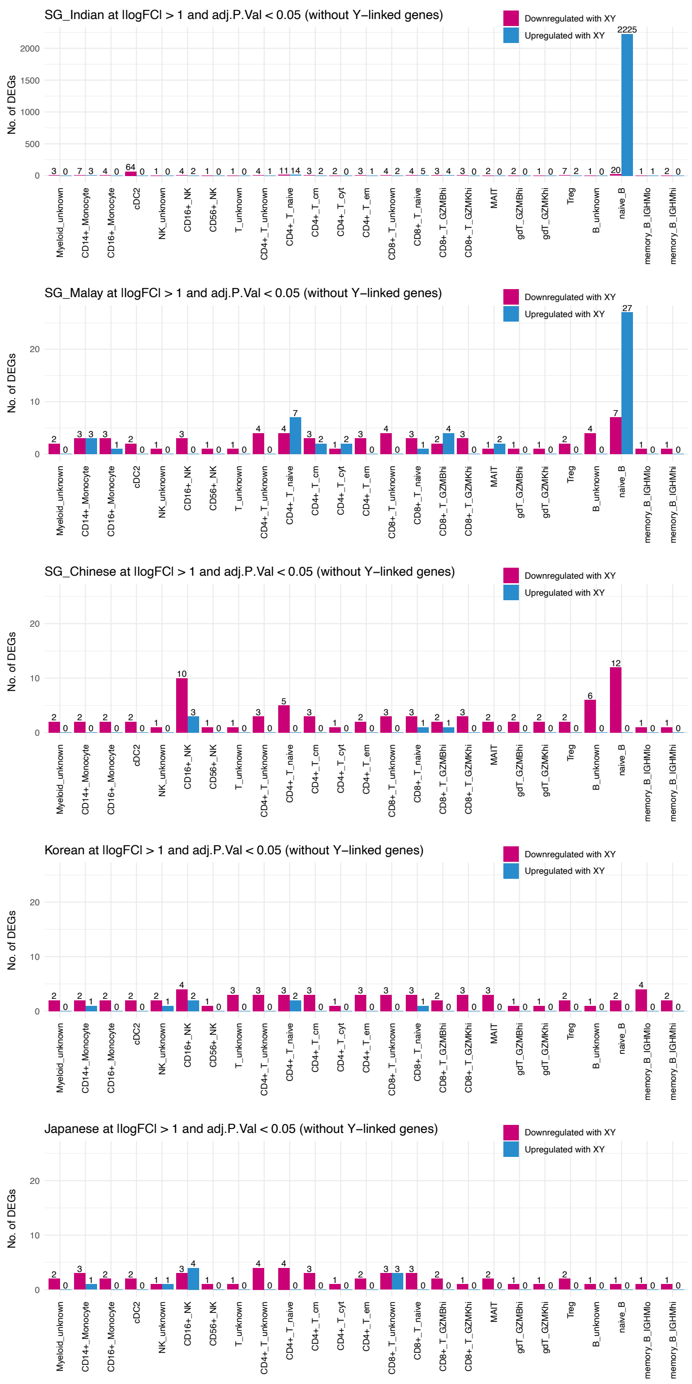


**Figure S24. Number of DEG associated with sex in each AIDA population.** The DEGs testing was using LMM per gene in each cell type of each individual (logcpm(Gene expression)$_{\text{cell type}}$ ~ Age + Sex + % Main Ancestry + (1|Library) at |logFC| > 1 and adj.P.Val < 0.05 of all DEGs without Y linked genes. “Main Ancestry” used in the model were Blue for AIDA Thailand, SG Chinese and SG Malay; Red for AIDA Japan and Korea; and Orange for SG India.

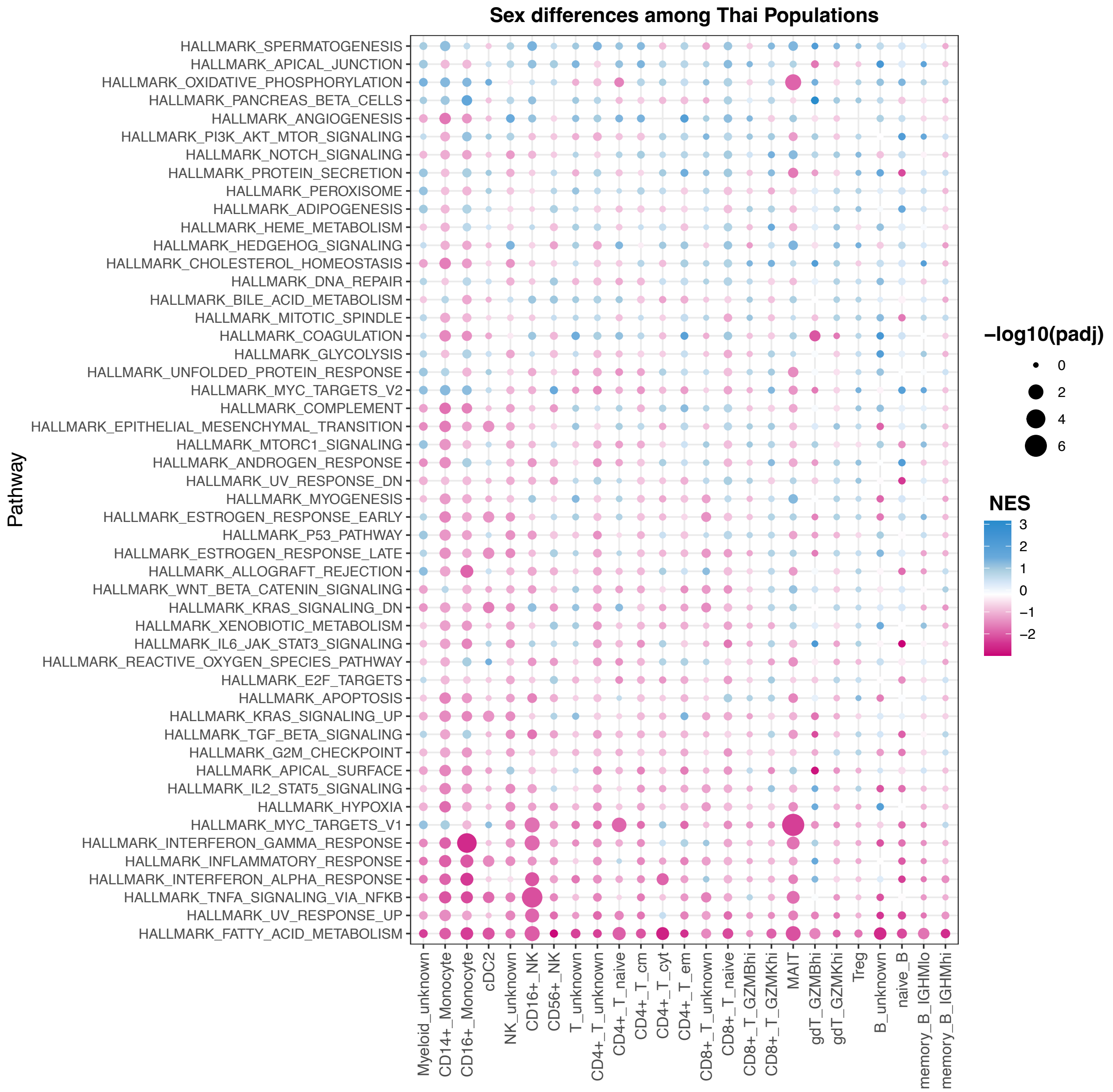


**Figure S25. Pathway enrichment of sex differences using GSEA analysis across all cell types in the Thai populations.**

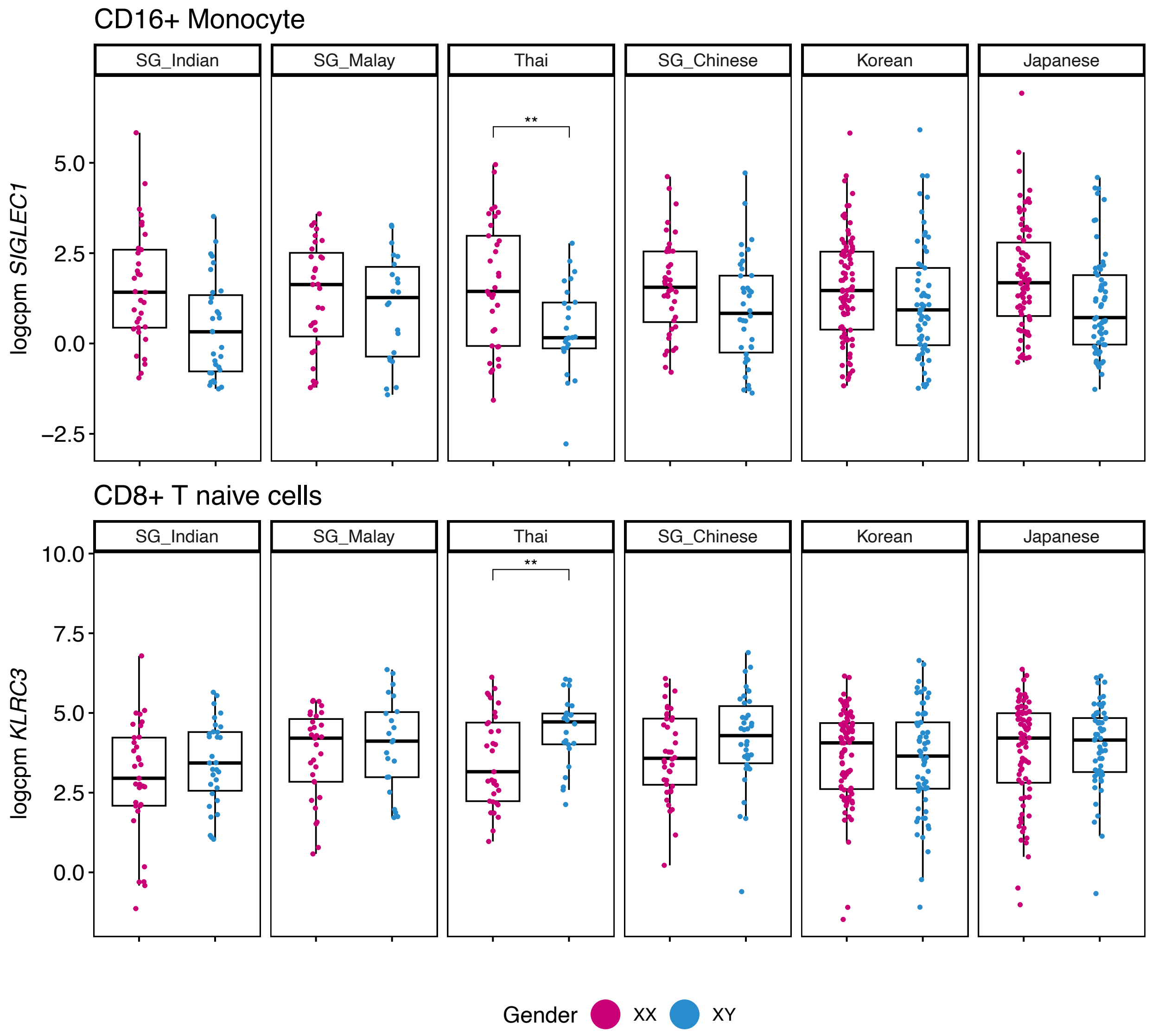


**Figure S26. Comparison of DEGs by sex differences in Thai and other AIDA populations.** The statistical testing was performed by LMM; ** $p \leq 0.01$ (logcpm(Gene expression)$_{\text{cell type}}$ ~ Age + Sex + % Main Ancestry + (1|Library). “Main Ancestry” used in the model were Blue for AIDA Thailand, SG Chinese and SG Malay; Red for AIDA Japan and Korea; and Orange for SG India.

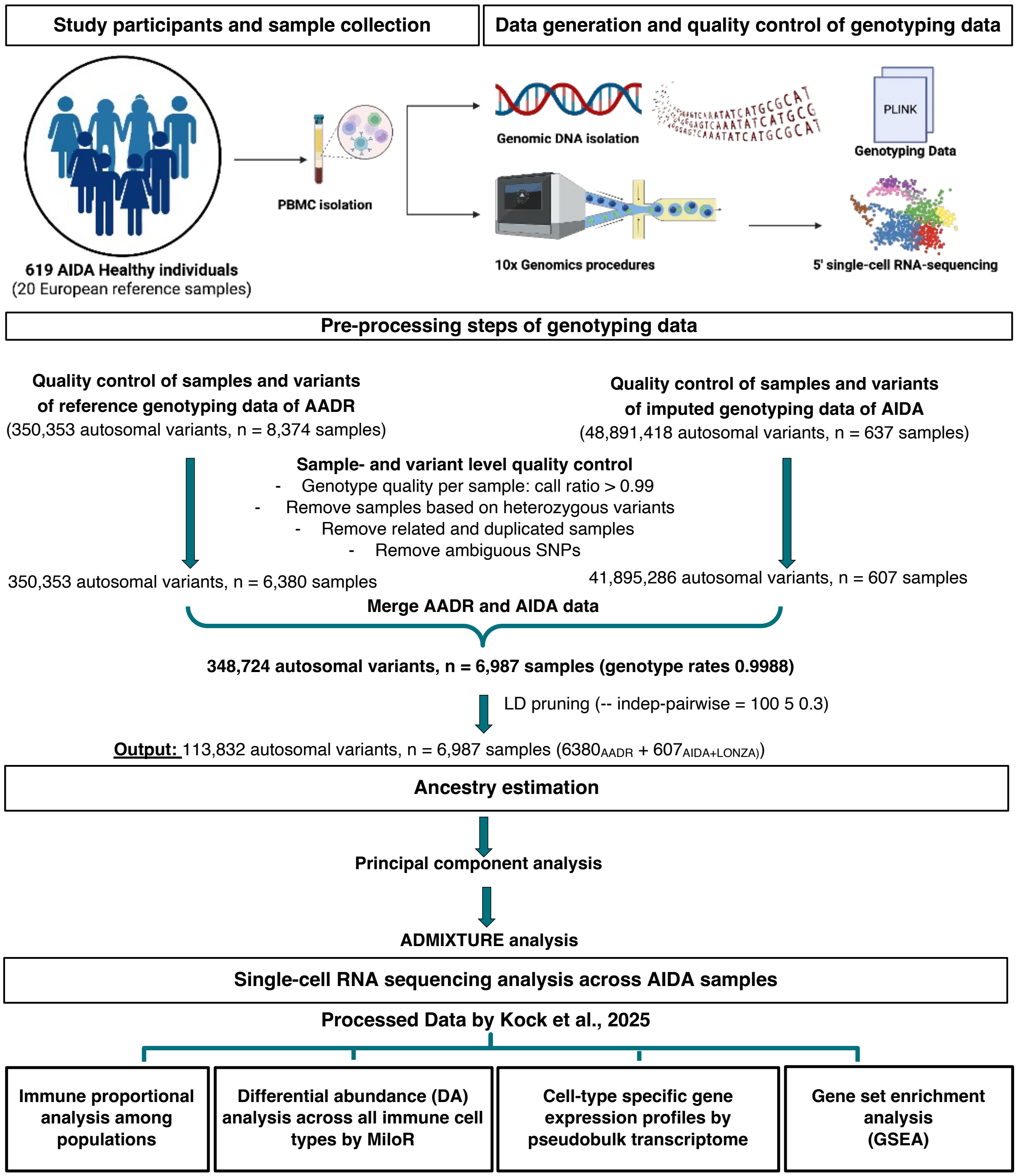


**Figure S27. Schematic workflow of all the analyses in this study**

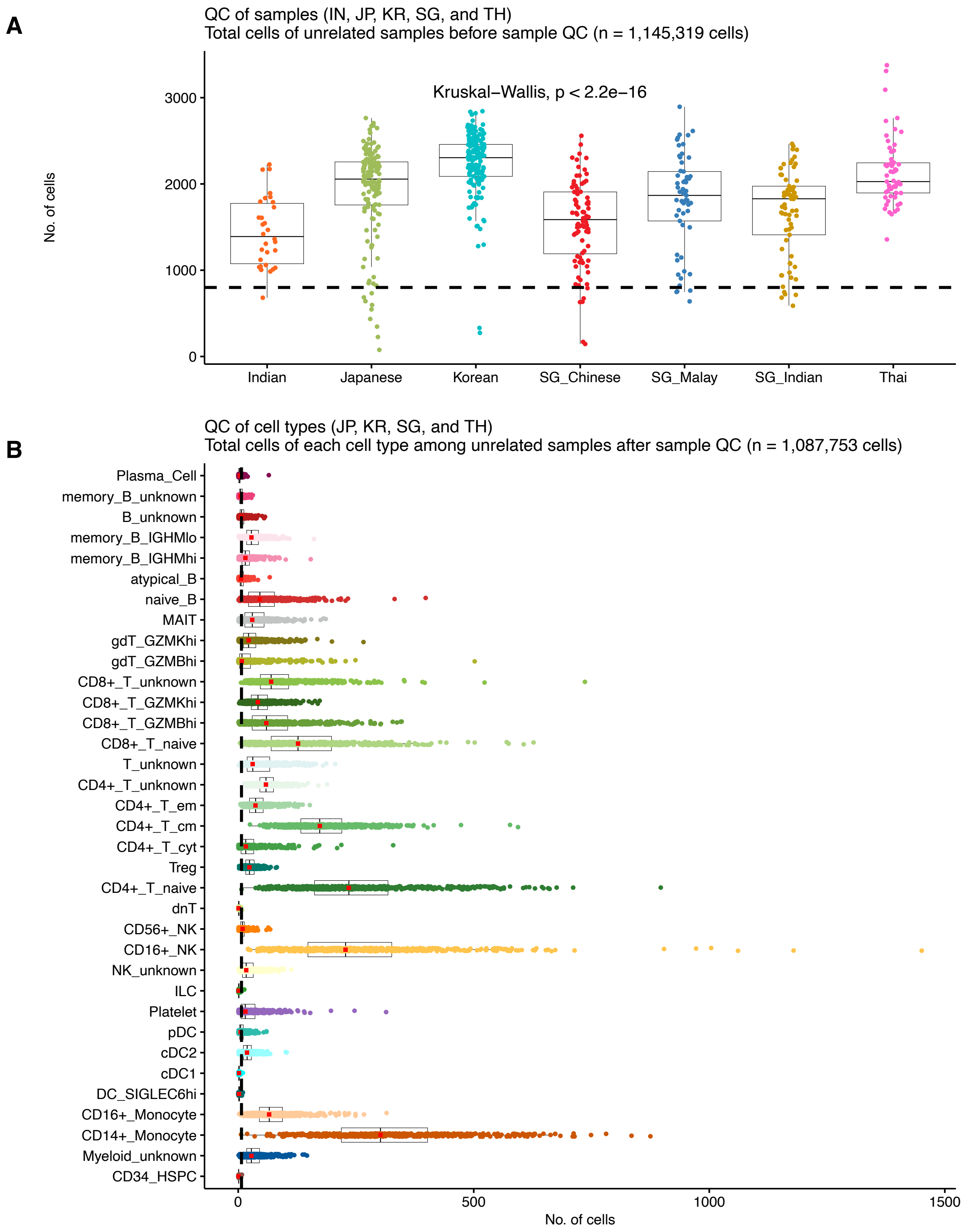


**Figure S28. Overall number of cells for cell proportional analysis.** QC steps were performed among **(A)** AIDA individuals and **(B)** cell subtypes. The black dashed line indicates the median cell count threshold used for QC.
